# Cross-species structure–mechanics prediction in the intervertebral disc

**DOI:** 10.64898/2026.10.30.763221

**Authors:** Jishizhan Chen, Alissa Parmenter, Aikta Sharma, Isla Henderson, Joseph Brunet, Stefan Zwingenberger, Deepak Bushan Raina, Theresa Urban, Hector Dejea, Sebastian Marussi, Eral Bele, Andrew A Pitsillides, Peter D Lee

**Affiliations:** Multiscale X-ray Imaging (MXI) Lab, Mechanical Engineering, UCL, WC1E 7JE, UK; Research Complex at Harwell, Rutherford Appleton Laboratory, Oxfordshire OX11 0FA, UK; European Synchrotron Radiation Facility, Grenoble, France; University Centre for Orthopaedic, University Hospital Carl Gustav Carus, Technical University Dresden, Dresden, Germany; Centre for Translational Bone, Joint and Soft Tissue Research, Technical University Dresden, Dresden, Germany; Faculty of Medicine, Department of Clinical Sciences, Orthopedics, Lund University, Sweden; Department of Comparative Biomedical Sciences, Royal Veterinary College, London NW1 0TU, UK

**Keywords:** Intervertebral disc, Annulus fibrosus, Fibre organisation, Synchrotron tomography, Digital volume correlation, Disc degeneration

## Abstract

Despite long-standing uncertainty regarding the suitability of animal intervertebral disc (IVD) models for the study of human IVD pathophysiology, few cross-species studies have integrated hierarchical structure with mechanical function. In this study, we present a multiscale structure–mechanics framework combining synchrotron computed tomography (sCT), fibre tracing, digital volume correlation (DVC) with structure-based mechanical inference and finite-element modelling. Rat, mini-pig and human IVDs (n = 3/species) were imaged by sCT, with rat and mini-pig discs additionally subjected to *in situ* uniaxial compression and DVC. A Structure Index (SI), integrating local structural anisotropy and fibre orientation, was developed to regionally pinpoint the location of mechanically vulnerable regions. Cross-species comparison showed scale-dependent structural conservation: gross disc and lamellar organisation retained common regional features, whereas fibre-bundle architecture exhibited greater species- and regional divergence. Despite these differences, lower SI was consistently associated with elevated deformation in rat and mini-pig discs across multiple strain measures. Structure–strain inverse correlations became tighter within progressively higher-strain domains, reaching Spearman ρ values of approximately −0.5 to −0.7 for several strain components. Synthetic fibre phantoms confirmed that SI reflects a coupling of fibre orientation and organisation, with additional sensitivity to fibre number in poorly organised configurations. In human discs with varying degrees of degeneration, SI revealed spatial heterogeneity and redistribution of structurally vulnerable regions. Fibril-informed homogenisation provided anisotropic material properties for organ-scale finite-element modelling that independently supported SI-based predictions. Overall, these findings establish a cross-species framework linking hierarchical collagen architecture to mechanical vulnerability and provide a basis for non-invasive, structure-driven biomechanical assessment of human IVD degeneration and mechanical risk.

## Introduction

Fibrous architecture is commonly found in various organs, such as the annulus fibrosus (AF) of the intervertebral disc (IVD)^1^, myocardial fibres in the heart^2^, collagen fibres in tendons^3^, and elastic fibres in the lung^4^. In these organs, fibre organisation provides structural anisotropy that enables tissues to withstand complex loading environments while maintaining functional integrity. Notably, collagen fibrils represent one of the most conserved structural elements across vertebrate species (∼10 nm to 500 nm in diameter, ∼50-fold variation)^5–7^, despite supporting organs that span several orders of magnitude in size (∼200 μm for a zebrafish embryo heart^8^ to ∼1 m in the blue whale ^9^, ∼5,000-fold variation) and differ in age^7^ and mechanical demand^10^. This suggests that the functional diversity of fibrous tissues arises not simply from fibril size itself, but also from architectural organisation across multiple scales. The structure-mechanics changes in fibres may determine how organs deform under physiological conditions or when they become pathologically degenerated and dysfunctional. In some cases, such as IVD degeneration, this represents key pathological features of disease^11–13^. Therefore, quantitatively linking fibre organisation to local mechanical behaviour is important for understanding degeneration mechanisms and identifying mechanically vulnerable regions. Spatial mapping of these regions may further refine the characterisation of IVD degeneration and reveal areas at risk of structural failure before macroscopic changes become apparent. However, such relationships remain difficult to resolve because tissue deformation emerges from heterogeneous fibre architectures operating across multiple scales^14–16^.

The IVD is a fibre-reinforced biological structure^17,18^. Its AF consists of lamellae of extracellular matrix with embedded collagen fibre bundles of alternating orientations. This hierarchical organisation allows the AF to resist multi-directional loading while stabilising the central nucleus pulposus^19^. Degeneration of the IVD disrupts fibre organisation which leads to altered mechanical behaviour, which is thought to be a main contributor to the development of lower back pain^20^. Current studies provide limited insight into the spatial heterogeneity of disc mechanics. To investigate the structural–mechanics correlation, animal models are widely used in IVD research. However, the suitability of different species remains debated^21,22^. Previous cross-species comparisons have largely focused on gross morphology^22^, biochemical composition^23,24^ or macroscopic mechanical properties^25,26^. These comparisons demonstrate that no single animal model reproduces all features of the human IVD. For example, rodent discs differ substantially from human discs in scale, whereas larger animal discs, including porcine models, still differ in loading environment, tissue composition, and geometry. Consequently, similarity in gross morphology or bulk mechanical behaviour does not imply similarity in the relationship between fibre organisation and deformation. Whether structure–mechanics relationships are conserved across species therefore remains unknown. Establishing this is important both for selecting appropriate animal models and for determining which observed mechanisms can be reasonably translated to human.

Recent advances in synchrotron computed tomography (sCT) have enabled visualisation of micro-scale soft tissues in whole organs^27^. When coupled with digital volume correlation (DVC), strain fields can be calculated under near-physiological loading conditions^28^. These developments provide an opportunity to investigate structure–mechanics relationships in fibrous tissues, however, extracting meaningful structural descriptors from complex fibre architectures remains a major challenge.

In this study, we use sCT and fibre tracing to reconstruct the three-dimensional (3D) fibre organisation of rat, mini-pig and human discs. Then, using these reconstructed images, we defined a Structure Index (SI) that quantifies lamellar anisotropy based upon structure tensor analysis and fibre orientation, and its subsequent combination with DVC-derived strain fields in animal discs, permits quantitative structure–strain correlation and the evaluation of its conservation across species. Finally, we applied the SI to static human disc datasets to predict mechanically vulnerable regions and investigated how these regions redistribute strain during disc degeneration, with validation using a fibril-informed finite element (FE) model. This structure–mechanics framework enables the prediction of tissue mechanics from static depictions of structural organisation, providing a potential route towards non-invasive risk assessment of disc degeneration and cross-species translation of structure–function relationships.

## Results

### Cross-species synchrotron imaging reveals conserved annulus fibrosus architectures with distinct morphologies

To determine whether structural organisation of the AF is conserved across species, we imaged intact rat, mini-pig, and human^29^ IVDs using phase-contrast sCT and, where feasible, under physiologically representative loads delivered in situ using a mechanical compression stage rig (**Fig. 1a**). The image reconstruction and fibre tracing was subsequently performed to characterise the 3D organisation of AF collagen bundles.

**Figure 1.**
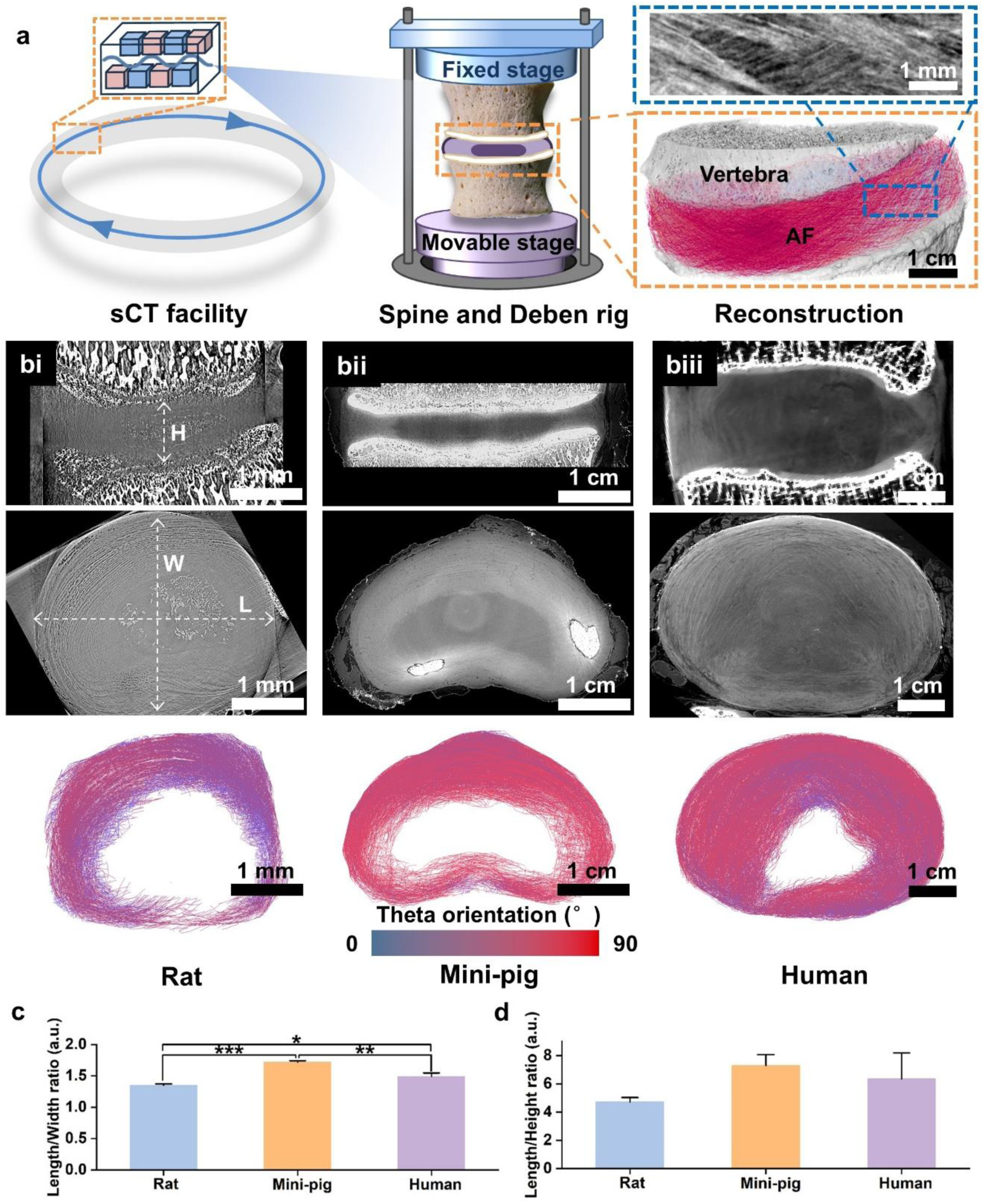
Experimental setup and overview of AF organisation across species. **(a)** sCT imaging setup with *in situ* mechanical loading using a custom Deben rig, illustrating sample mounting between fixed and movable stages and image acquisition, reconstruction, and fibre visualisation workflow (mini-pig lumbar L4/5 as an example here); **(b)** Cross-species comparison of lumbar L4/5 IVDs in **(bi)** rat, **(bii)** mini-pig, and **(biii)** human. Top and middle rows show sagittal and transverse sCT images, and the bottom row shows corresponding colour-coded fibre theta (θ) orientation, the polar angle between a fibre and the z axis; **(c)** Length-to-width ratio and **(d)** length-to-height ratio of the IVDs across species. Statistical comparisons were performed using one-way ANOVA with Tukey’s post hoc comparison. *: P < 0.05; **: P < 0.01; ***: P < 0.001.

Representative sagittal (top row, **Fig. 1b**) and transverse (middle row, **Fig. 1b**) sCT reconstructions demonstrated substantial differences in overall disc size between rat (L2/3, L4/5, and L5/6), mini-pig (L4/5), and human (L4/5) IVDs. Rat IVDs (length = ∼4 mm) were approximately one order of magnitude smaller than mini-pig (length = ∼30 mm) and human IVDs (length = ∼60 mm), while human IVDs were about twice the size of mini-pig IVDs. Gross morphology comparison revealed a conserved ring-like AF surrounding the central nucleus pulposus together with broadly preserved fibre θ orientation patterns across species (full IVD mean θ = 54° – 63°, P > 0.05, bottom row, **Fig. 1b**). This overall architecture aligns with the known fibre-reinforced organisation of mammalian IVDs. However, substantial species-dependent differences in morphology and AF organisation were also observed. The mini-pig IVD exhibited a horseshoe-shaped morphology, whereas rat and human discs retained a more oval geometry. IVD length-to-width ratios were 1.35 ± 0.02, 1.72 ± 0.02, and 1.49 ± 0.05 in rat, mini-pig, and human IVDs, respectively (**Fig. 1c** and **Table 1**). The mini-pig IVD exhibited a significantly greater length-to-width ratio than both the rat (P < 0.001) and human IVDs (P < 0.01), while a smaller but statistically significant difference was also observed between rat and human IVDs (P < 0.05). Mini-pig IVDs showed the highest length-to-height ratio (7.29), followed by human (6.35) and rat (4.73) IVDs. Thus mini-pig IVDs visually appeared to have a relatively narrower intervertebral space, although differences in length-to-height ratio among species were not statistically significant (P > 0.05; **Fig. 1d** and **Table 1**).

**Table 1.**
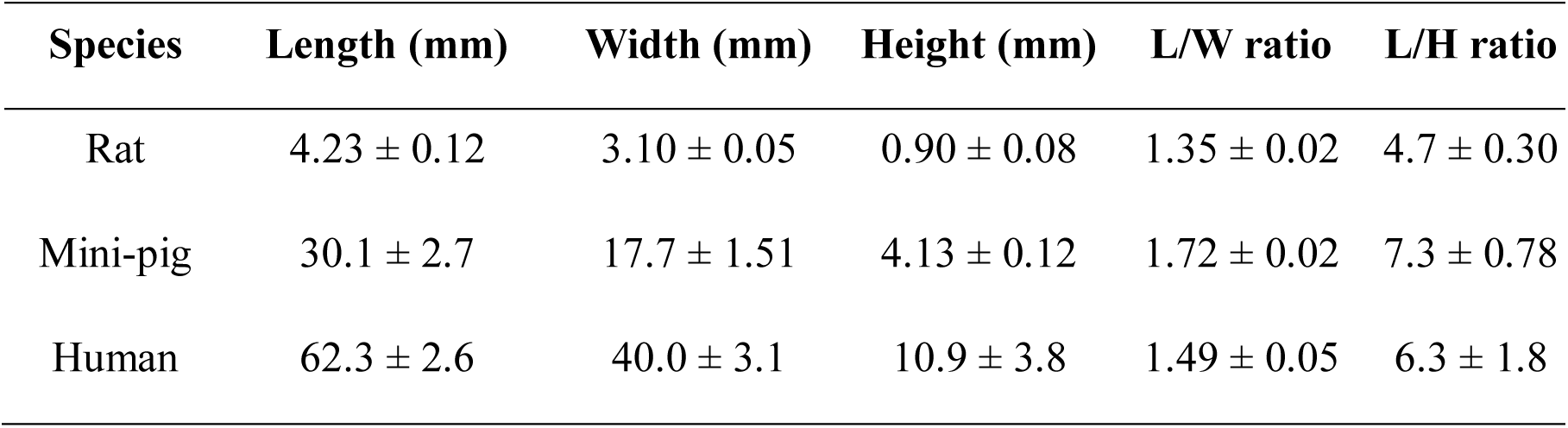
Gross comparisons of rat (n=3, L2/3, L4/5, and L5/6), mini-pig (n=3, L4/5), and human (n=3, L4/5) IVD dimensions.

| **Species** | **Length (mm)** | **Width (mm)** | **Height (mm)** | **L/W ratio** | **L/H ratio** |
|---|---|---|---|---|---|
| Rat | 4.23 ± 0.12 | 3.10 ± 0.05 | 0.90 ± 0.08 | 1.35 ± 0.02 | 4.7 ± 0.30 |
| Mini-pig | 30.1 ± 2.7 | 17.7 ± 1.51 | 4.13 ± 0.12 | 1.72 ± 0.02 | 7.3 ± 0.78 |
| Human | 62.3 ± 2.6 | 40.0 ± 3.1 | 10.9 ± 3.8 | 1.49 ± 0.05 | 6.3 ± 1.8 |

These findings indicate that although the overall AF architecture is conserved across species, substantial divergence exists in disc size, morphology, and aspect ratio, raising the question of whether underlying structure–mechanics relationships are similarly conserved.

### Lamellar organisation is regionally conserved but spatially heterogeneous across species

To determine whether conservation of gross AF architecture also extends to the lamellar scale, we quantified regional and cross-species variations in AF lamellar organisation. Anterior and posterior–lateral AF regions were extracted for analysis across the disc height (**Fig. 2ai–aiii**). Within each XY plane, pixel-wise radial sampling paths were generated across the AF from the outer towards the inner surface, enabling spatially resolved quantification of lamellar organisation (**Fig. 2b**). Line-profile analysis of grayscale intensity along individual radial paths demonstrated periodic oscillations corresponding to individual lamellae, enabling quantitative lamellar counting (**Fig. 2c**).

**Figure 2.**
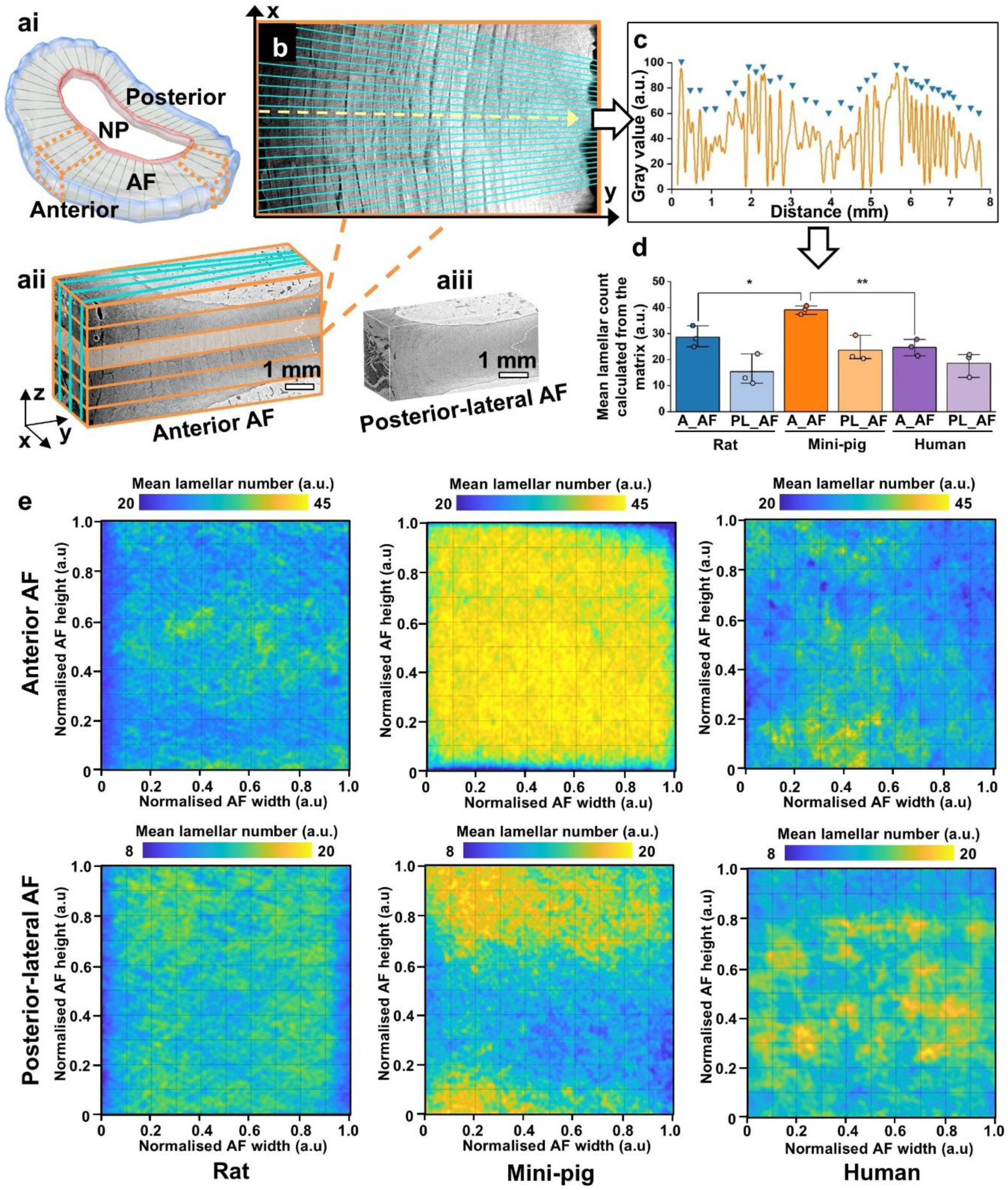
Cross-species spatial mapping of AF lamellar organisation. **(ai)** Schematic showing the anterior and posterior-lateral AF regions selected for analysis. Representative sCT subvolumes from the **(aii)** anterior and **(aiii)** posterior-lateral AF, respectively. Orange lines indicate sequential XY planes analysed through the AF height, whereas cyan lines indicate pixel-wise radial sampling paths along the *x* direction from the outer to the inner AF, used to quantify the number of lamellae intersected along each path. Both lines are displayed at reduced density for visualisation; **(b)** Representative XY plane illustrating pixel-wise radial sampling from the outer to the inner AF across the lamellar structure; **(c)** Representative intensity profile along a sampling path. Triangles indicate detected peaks correspond to individual lamellae; **(d)** Mean lamellar number in the anterior (A_AF) and posterior-lateral (PL_AF) AF of rat, mini-pig, and human IVDs, calculated as the mean ± SD of all values across the corresponding width–height lamellar number matrix for each specimen. Statistical comparisons were performed using one-way ANOVA with Tukey’s post hoc comparison. *: P < 0.05; **: P < 0.01; ***: P < 0.001; **(e)** Spatial maps of local lamellar number across normalised AF width and height in the anterior and posterior-lateral AF of rat, mini-pig, and human IVDs.

The anterior AF contained 1.3–1.9-fold more lamellae than the posterior–lateral AF in all species, with the greatest lamellar counts observed in mini-pig discs. Mean lamellar number across the complete width–height matrix was 39.1 ± 1.3 in the mini-pig anterior AF, compared with 28.7 ± 3.3 in rat (P < 0.05) and 24.7 ± 2.6 in human discs (P < 0.01) (**Fig. 2d**). Although posterior–lateral AF consistently exhibited lower mean lamellar numbers than the corresponding anterior region, no significant differences were observed (P > 0.05). We also observed a depth-dependant change of lamellar thickness. Using a representative mini-pig disc as an example, there was a progressive decrease in lamellar thickness from the outer to inner AF, from approximately 300–400 μm to ∼80 μm in the anterior and from approximately 100–150 μm to ∼40 μm in the posterior–lateral (**Fig. S1**). Across species and at equivalent relative depths, anterior AF lamellae were approximately 2–3-fold thicker than posterior–lateral lamellae, demonstrating conserved regional gradients in lamellar architecture. As the total number of lamellae varied among samples, no statistical comparisons were performed for AF thickness.

Spatial analysis further showed that lamellar number was non-uniform across both AF width and height (**Fig. 2e** and **Fig. S2**). In the anterior AF, mini-pig IVDs maintained relatively high lamellar numbers across most of the normalised width–height matrix (approximately 40–45), accompanied by comparatively low inter-sample variability over the central region (**Fig. S2a**). Rat and human anterior AFs showed broader spatial heterogeneity, with local lamellar numbers predominantly spanning approximately 25–32 and regions of increased variability (SD up to ∼9), particularly across the lower half of the rat AF and more broadly distributed regions in the human AF (**Fig. S2a**). Posterior–lateral AF exhibited lower local lamellar numbers overall (approximately 8–20). Mini-pig IVDs showed greater lamellar number in the upper and lower regions, whereas rat and human IVDs exhibited discrete localised regions of increased lamellar number over the entire matrix (**Fig. S2b**). Inter-sample variability in the posterior–lateral AF remained generally lower than in the anterior AF (SD predominantly within 0–6). Notably, the human IVD specimens exhibited varying degrees of degeneration; therefore, the spatial heterogeneity observed in the human AF likely reflects both inter-species differences and degeneration-associated structural remodelling. Overall, these spatial maps demonstrate that differences extend beyond mean lamellar number to the regional variability of lamellar organisation across the AF.

In addition to these expected lamellar features, sCT imaging further revealed frequent branching and extension of adjacent lamellae, forming continuous interconnected structures rather than isolated concentric sheets (**Fig. S3** and **Supplementary Videos 1–2**). These observations suggest that AF lamellae undergo substantial interweaving and structural reorganisation across the tissue depth. This interconnected organisation has been impossible to visualise directly using conventional imaging approaches and represents a previously underappreciated architectural feature of intact whole-disc AF organisation.

These similarities and differences observed at both the gross morphological and lamellar levels motivated further investigation into whether finer fibre-bundle architecture shows similar conservation or greater structural divergence across species.

### Fibre-bundle organisation diverges across species despite conserved lamellar organisation

Given the broadly conserved lamellar organisation across species, we next investigated whether this conservation also extends to the finer fibre-bundle scale using fibre tracing. In contrast to the relatively conserved lamellar organisation, AF fibre-bundle architecture exhibited substantial regional and species-dependent variation. Fibre tracing in the anterior AF revealed coherent fibre bundles spanning the outer, middle, and inner AF regions in all species (**Fig. 3a**). Consistent with the known functional organisation of the AF, fibres in the outer AF were more uniformly aligned, whereas the inner AF displayed progressively greater orientation dispersion.

**Figure 3.**
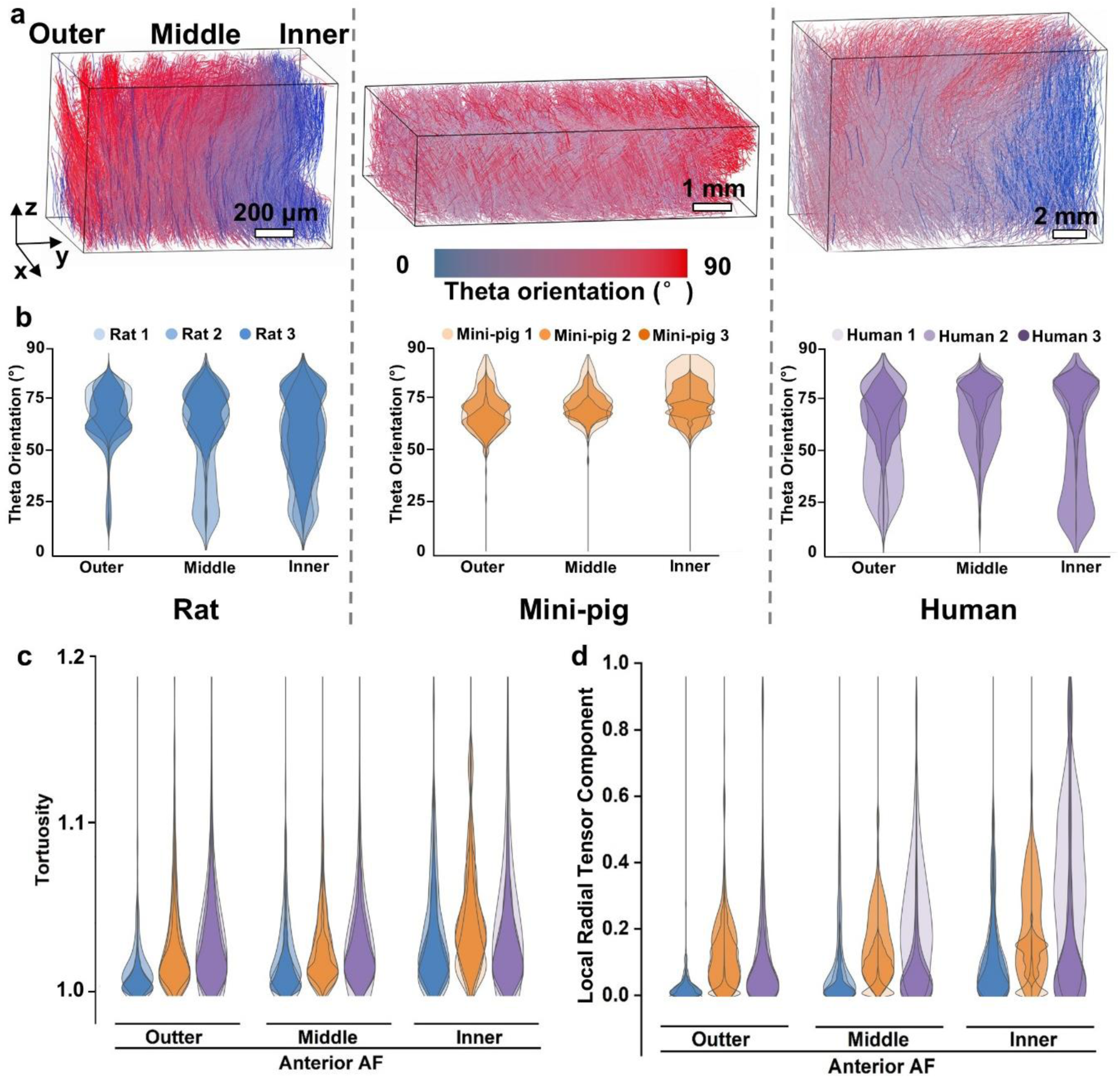
Quantitative analysis of fibre organisation in the anterior AF. **(a)** Representative fibre architectures across outer, middle, and inner regions of the anterior AF. Fibre orientation is colour-coded by θ angle; **(b)** Distribution of fibre θ angle across species (rat, mini-pig, and human) and radial regions; **(c)** Fibre tortuosity across outer, middle, and inner regions for each species; **(d)** Local radial tensor component across radial regions and species, quantifying fibre orientation relative to the nucleus pulposus.

However, quantitative orientation analysis revealed species-dependent divergence in these radial organisational patterns. In rat and human discs, θ angle showed a decreasing trend from the outer to inner AF, indicating progressively more perpendicular fibre orientation. θ angles were comparable between rat and human discs in both the outer (65° ± 10° vs 62° ± 15°, P>0.05) and inner AF (49° ± 19° vs 57° ± 25°, P>0.05). In contrast, mini-pig discs showed an increasing trend from the outer to inner AF (from 65° ± 9° to 72° ± 10°). While outer AF θ angles were comparable between mini-pig and both rat and human discs (both P > 0.05), inner AF θ angles were higher in mini-pig than rat discs (P < 0.01), but comparable between mini-pig and human discs (P > 0.05, **Fig. 3b**). Fibre tortuosity increased from outer to inner AF in all species and was highest in human (up to 1.043±0.094) and lowest in rat discs (up to 1.033±0.036, **Fig. 3c**). The local radial tensor component, reflecting fibre orientation relative to the nucleus pulposus, increased radially in rat and mini-pig discs but varied substantially in human discs (**Fig. 3d**). These demonstrate that although overall AF lamellar organisation is conserved, fibre-bundle architecture exhibits considerable species-specific divergence at finer structural scales.

Posterior–lateral AF fibre bundles exhibited greater spatial heterogeneity across the AF thickness than the anterior AF (**Fig. 4a**), consistent with previously reported regional heterogeneity in AF fibre mechanics and strain distributions^28,30^. In rat and human discs, θ angle also showed a decreasing trend from the outer to inner posterior–lateral AF. θ angles were comparable between rat and human discs in both the outer (55° ± 20° vs 60° ± 14°, P > 0.05) and inner AF (39° ± 18° vs 52° ± 25°, P > 0.05). In contrast, mini-pig discs showed an increasing trend from the outer to inner AF (from 64° ± 11° to 73° ± 8°). While outer AF θ angles were comparable between mini-pig and both rat and human discs (both P > 0.05), inner AF θ angles were higher in mini-pig than rat discs (P < 0.01), but comparable between mini-pig and human discs (P > 0.05, **Fig. 4b**). Despite the broad similarity between the anterior and posterior–lateral AF, θ-angle dispersion in the posterior–lateral AF was broader than in the anterior AF at corresponding depths, indicating weaker radial organisation and reduced regional gradients. Posterior–lateral fibre tortuosity also showed substantial inter-species heterogeneity compared with the anterior AF (**Fig. 4c**). In contrast to the anterior AF, the local radial tensor component decreased towards the nucleus pulposus in all species (**Fig. 4d**).

**Figure 4.**
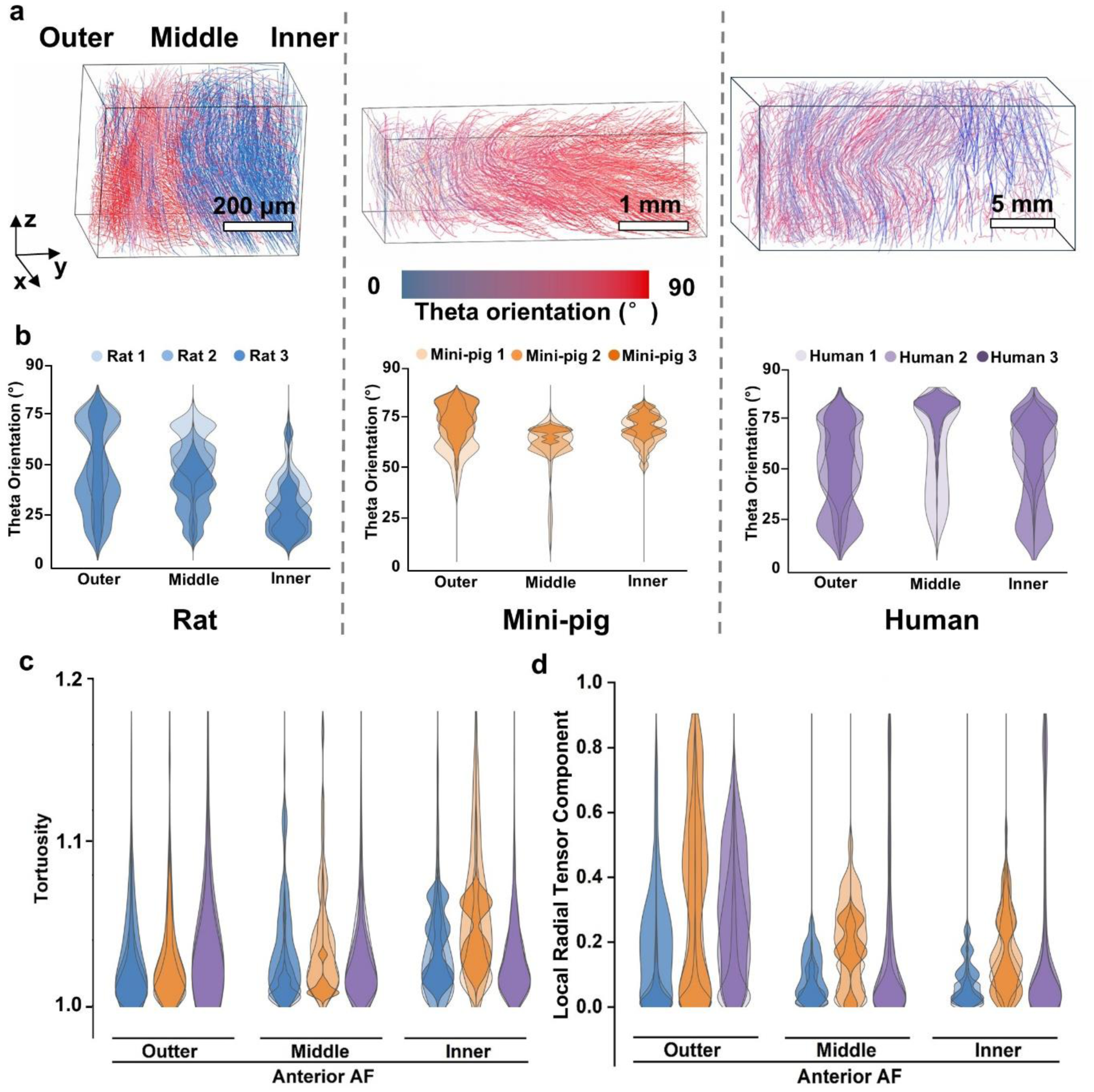
Quantitative analysis of fibre organisation in the posterior–lateral AF. **(a)** Representative fibre architectures across outer, middle, and inner regions of the posterior–lateral AF at different spatial scales. Fibre orientation is colour-coded by θ angle; **(b)** Distribution of fibre θ angle across species (rat, mini-pig, and human) and radial regions; **(c)** Fibre tortuosity across outer, middle, and inner regions for each species; **(d)** Local radial tensor component across radial regions and species, quantifying fibre orientation relative to the nucleus pulposus.

These findings indicate that structural conservation observed at the lamellar level does not fully extend to the fibre-bundle scale. Instead, fibre-bundle organisation exhibits substantial species-and region-specific divergence. This raised the question of whether mechanically relevant structure–strain relationships remain conserved despite these architectural differences.

### Structure Index identifies mechanically vulnerable regions across species

To investigate the structure–strain relationships across species, we developed a SI framework that integrates local structural anisotropy and fibre orientation to identify mechanically vulnerable regions. Spatial correlations between SI and DVC-derived strain fields were evaluated in rat and mini-pig IVDs. In both species, regions with lower SI values were associated with higher magnitudes of first and third principal strains, whereas regions with higher SI corresponded to lower strain magnitudes (**Fig. 5a**). Radial cross-sectional analysis further demonstrated that concentrated strains were predominantly located within the inner two-thirds of the anterior AF but within the outer one-third of the posterior–lateral AF in both rat and mini-pig discs (**Fig. 5b**). These conserved spatial patterns suggest that structurally vulnerable regions are similarly organised across species despite differences in fibre-bundle architecture.

**Figure 5.**
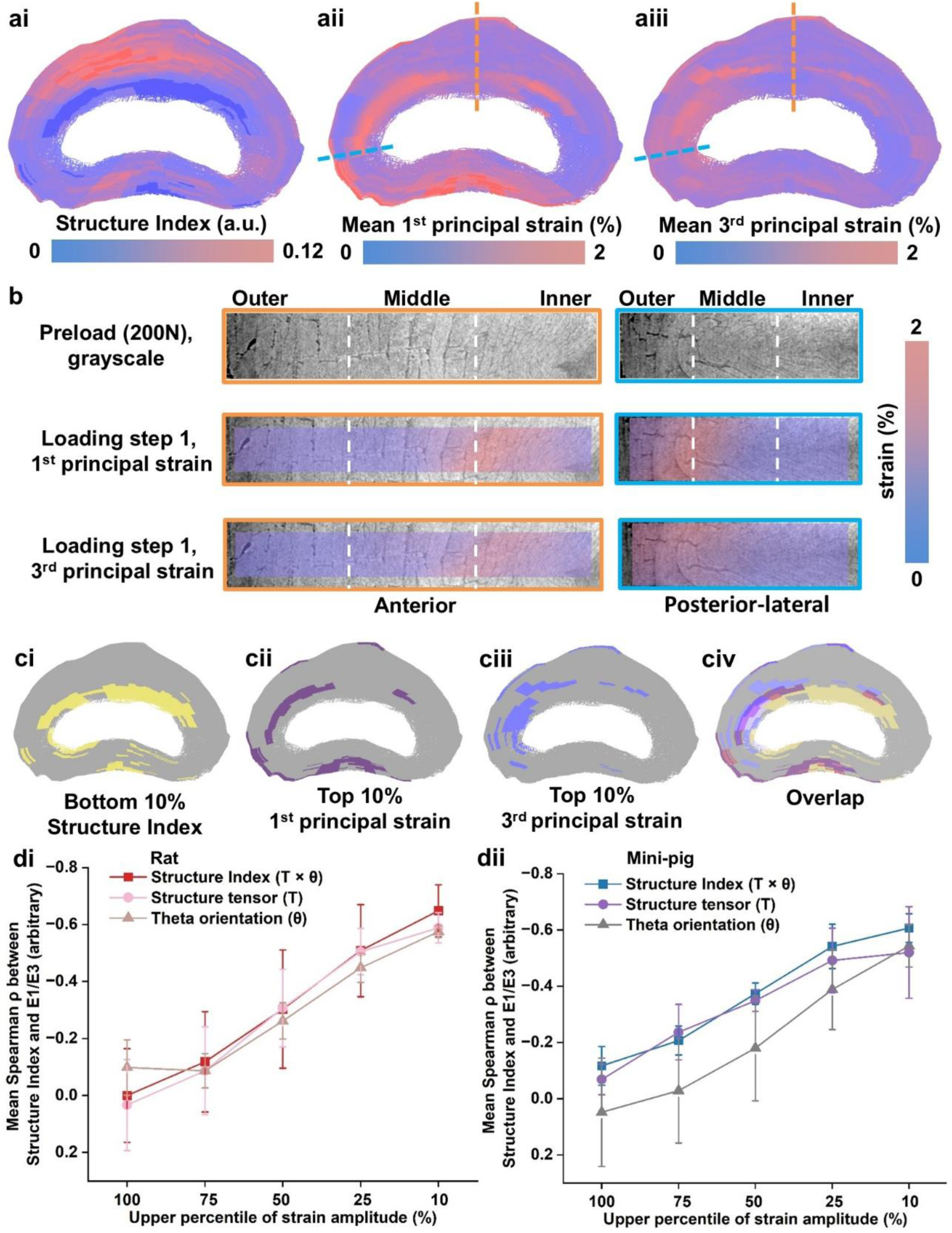
Spatial correlation between Structure Index and strain distribution in rat and mini-pig. **(a)** Spatial correspondence between structural organisation and mechanical response in the AF; **(ai)** SI distribution; **(aii)** Mean first principal strain distribution; **(aiii)** Mean third principal strain distribution; **(b)** Regional analysis of anterior and posterior–lateral AF under loading. Grayscale images (top) and corresponding first and third principal strain fields (middle and bottom); **(c)** Spatial overlap between structurally predicted vulnerable regions and high-strain regions; **(ci)** Bottom 10% SI; **(cii)** Top 10% first principal strain; **(ciii)** Top 10% third principal strain; **(civ)** Overlap between low SI and high-strain regions; **(d)** Quantification of structure–strain correlation across species: **(di)** Rat; **(dii)** Mini-pig.

Interestingly, only 3–10% spatial overlap was observed between low-SI regions (bottom 10%) and the highest-strain regions (top 10%; **Fig. 5c**) and was consistently observed across multiple strain measures, including principal, Von Mises, radial, circumferential and shear strains in both rat and mini-pig (**Fig. S4a** and **S5a**). This finding indicates that SI does not directly map instantaneous strain magnitude point-by-point, but instead identifies mechanically vulnerable structural configurations where large deformations preferentially initiate.

Spearman correlation analysis further demonstrated that structure–strain correlations increased progressively from full-field analysis to the highest-strain regions, reaching up to ρ ≈ −0.6 in the top 10% first and third principal strain domains (**Fig. 5d**). This trend was also observed for Von Mises, circumferential and shear strains, although the magnitude of the relationship varied between strain components and species (ρ ≈ −0.1–−0.7, **Fig. S4b** and **S5b**). Interestingly, the individual SI components, including structure tensor anisotropy and fibre θ orientation, exhibited similar trends but the combined SI consistently achieved the strongest inverse correlations within high-strain regions. These findings demonstrate that integrating anisotropy and fibre orientation captures mechanically relevant architectural information more effectively than either parameter independently.

To further investigate the origin of this combined effect, we generated a series of synthetic lamellar fibre phantoms to isolate the respective contributions of structural anisotropy, fibre orientation, and fibre number (**Supplementary Methods**). The simulations demonstrated that SI increased only when strong lamellar organisation was combined with larger θ angles (**Fig. S6**), establishing that SI reflects a coupled structural–orientational feature of fibre organisation rather than a simple representation of either component alone. Fibre number also influenced SI (**Fig. S7**); when the number of fibres increased from 3 to 15, the mean SI across all θ angles decreased predominantly in phantoms with low-to-moderate organisation, by 27.0%, 30.1%, and 27.2% at Order = 0, 0.25, and 0.5, respectively. Across all individual θ and Order combinations, the largest reduction was 36.2% at θ = 67.5° and Order = 0.25. In more organised phantoms, the mean SI decreased by only 6.7% and 3.1% at Order = 0.75 and 1, respectively, with the smallest reduction of 1.4% observed at θ = 22.5° and Order = 0.75. These results indicate that SI is primarily governed by fibre orientation and organisation, although a low number of fibres may lead to overestimation in disorganised structures.

The preferential structure–strain correlation observed in high-strain domains further suggests that SI is relatively insensitive to mechanically stable low-strain regions but selectively identifies structurally vulnerable architectures associated with concentrated deformation. Importantly, the medium-to-high correlations observed consistently in both rat and mini-pig discs indicate that these structure–mechanics relationships remain conserved across species despite substantial architectural divergence at the fibre-bundle scale. This conservation supports the subsequent application of SI to human discs for structure-based prediction of mechanically vulnerable regions without direct mechanical testing.

### Multiscale homogenisation links fibril architecture to organ-scale mechanical behaviour in human discs

Having established structure–mechanics relationships in rat and mini-pig IVDs, we examined whether this approach could be extended to human IVDs. As direct DVC measurements were not possible in human specimens, we established a multiscale homogenisation framework to generate material properties for FE modelling and subsequent validation of SI-predicted mechanically vulnerable regions. Transmission electron microscopy imaging and segmentation enabled identification of individual collagen fibrils together with extraction of their geometrical features^31^. Fibril diameters exhibited a broad distribution ranging from approximately 10 to 250 nm, with a dominant population between 40 and 120 nm. The distribution was positively skewed, indicating a higher frequency of small-diameter fibrils with fewer large fibrils.

The spatial packing of fibrils was further characterised by calculating nearest-neighbour distances between fibril centres (**Fig. 6a**). Inter-fibril distances ranged from ∼60 to 215 nm, with a mean value of ∼130 nm. The spacing distribution followed a near-normal profile (Kolmogorov–Smirnov test, P > 0.05), indicating that fibril organisation could be approximated statistically using a Gaussian distribution. These experimentally derived descriptors were subsequently used to constrain stochastic generation of representative fibrillar microstructures.

**Figure 6.**
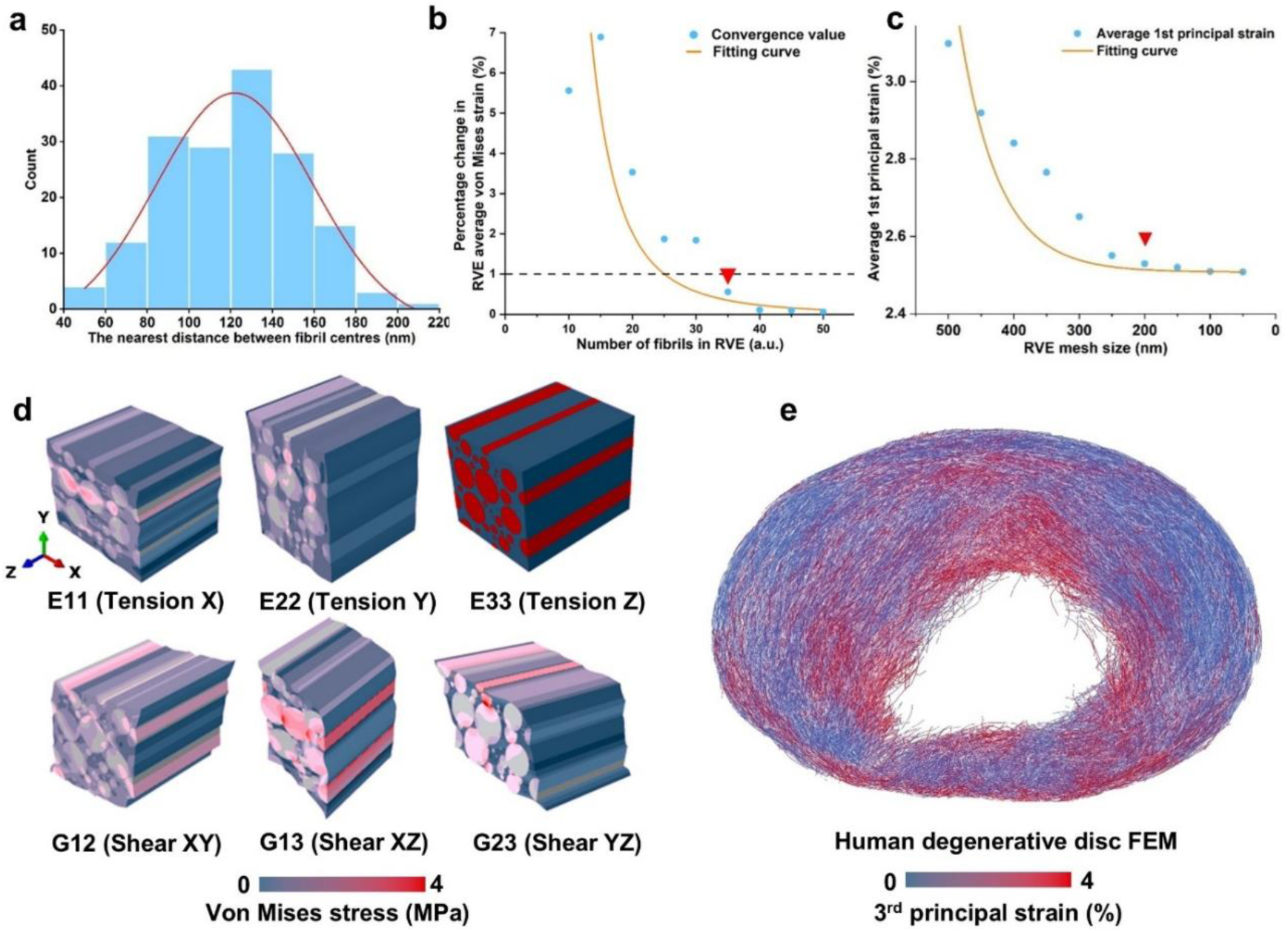
Multiscale translation from fibril organisation to tissue mechanics. **(a)** Nearest-neighbour distance distribution of fibril centres; **(b)** RVE size convergence characterised by the percentage change in volume-averaged von Mises strain with increasing fibril number. The dashed line indicates the 1% convergence threshold, and the red marker indicates the selected fibril number for RVE; **(c)** Stabilisation of the average first principal strain with decreasing RVE mesh size. The red marker indicates the selected RVE mesh size; **(d)** Homogenised stress fields under orthogonal loading conditions used to derive effective elastic engineering constants; **(e)** Organ-scale FEM strain distribution predicted using the RVE-derived engineering constants.

To determine the minimum representative microstructural size required for stable mechanical homogenisation, we evaluated the influence of fibril number on the homogenised mechanical response. As fibril number increased from 5 to 50, variability in volume-averaged von Mises strain progressively decreased. At low fibril numbers (≤15), relative variation exceeded 10%, indicating strong dependence on local geometrical arrangement. Variability decreased to ∼2–3% at 25–30 fibrils and further reduced below 1% at 35 fibrils (**Fig. 6b**), after which additional fibrils produced negligible changes (<0.5%). This convergence behaviour established a minimum representative volume element (RVE) size required to capture stable fibril-scale mechanical behaviour. Similarly, mesh convergence analysis demonstrated stabilisation at a mesh size of 200 nm with relative variation below 1% (**Fig. 6c**). The converged RVE was subsequently used for multiscale homogenisation.

The homogenised engineering constants extracted from the converged RVE demonstrated pronounced orthotropic behaviour (**Fig. 6d** and **Table 2**). This anisotropy emerged directly from the experimentally measured fibril-reinforced architecture, in which load transfer is governed by fibril alignment, packing, and connectivity. Resulting homogenised engineering constants were subsequently assigned to the AF within the organ-scale model (**Fig. 6e**), enabling mechanical validation of SI-predicted vulnerable regions in human IVD. Unlike conventional FE that relies on assumed or phenomenological material properties, this framework propagates experimentally measured fibril organisation directly into organ-scale mechanics, providing a physically grounded basis for structure-driven mechanical inference in human discs. Methodologies for human AF homogenisation are provided in **Supplementary Methods**.

**Table 2.**
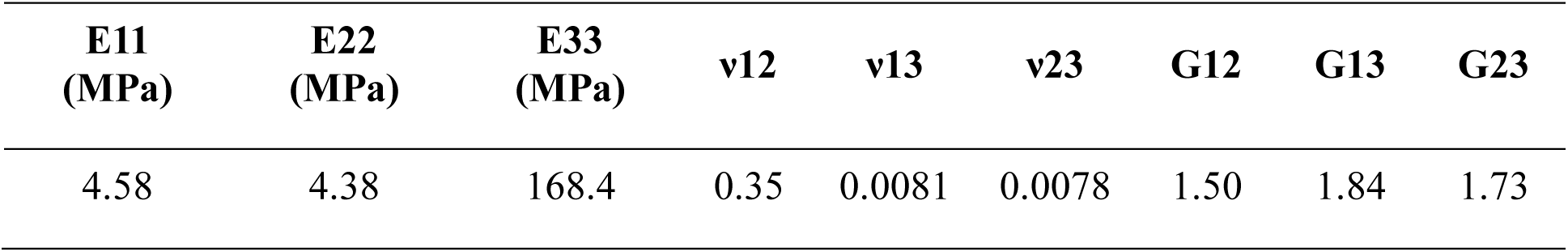
Homogenised material properties of the RVE.

### Degeneration alters spatial distribution of mechanically vulnerable human IVD regions

Having established a multiscale FEM framework for validating structure-driven mechanical vulnerability, we next investigated how degeneration affects the spatial redistribution of mechanically vulnerable regions in the human IVD. The SI framework was therefore appliedto static human disc datasets (**Fig. 7ai**)^29^ to predict mechanically vulnerable regions without direct mechanical testing. In the sCT image of healthy human AF (L1/2), the lamellae maintained ordered architecture and clear boundary between the nucleus pulposus and AF (**Fig. 7aii**). The spatial distribution of SI followed a pattern broadly consistent with that observed in rat and mini-pig discs. Low-SI regions were primarily located within the inner two-thirds of the anterior AF and the outer one-third of the posterior–lateral AF, resulting in a largely symmetric distribution across the disc (**Fig. 7aii**). This conserved organisation supports the cross-species generalisability of the previously identified structure–strain relationships.

**Figure 7.**
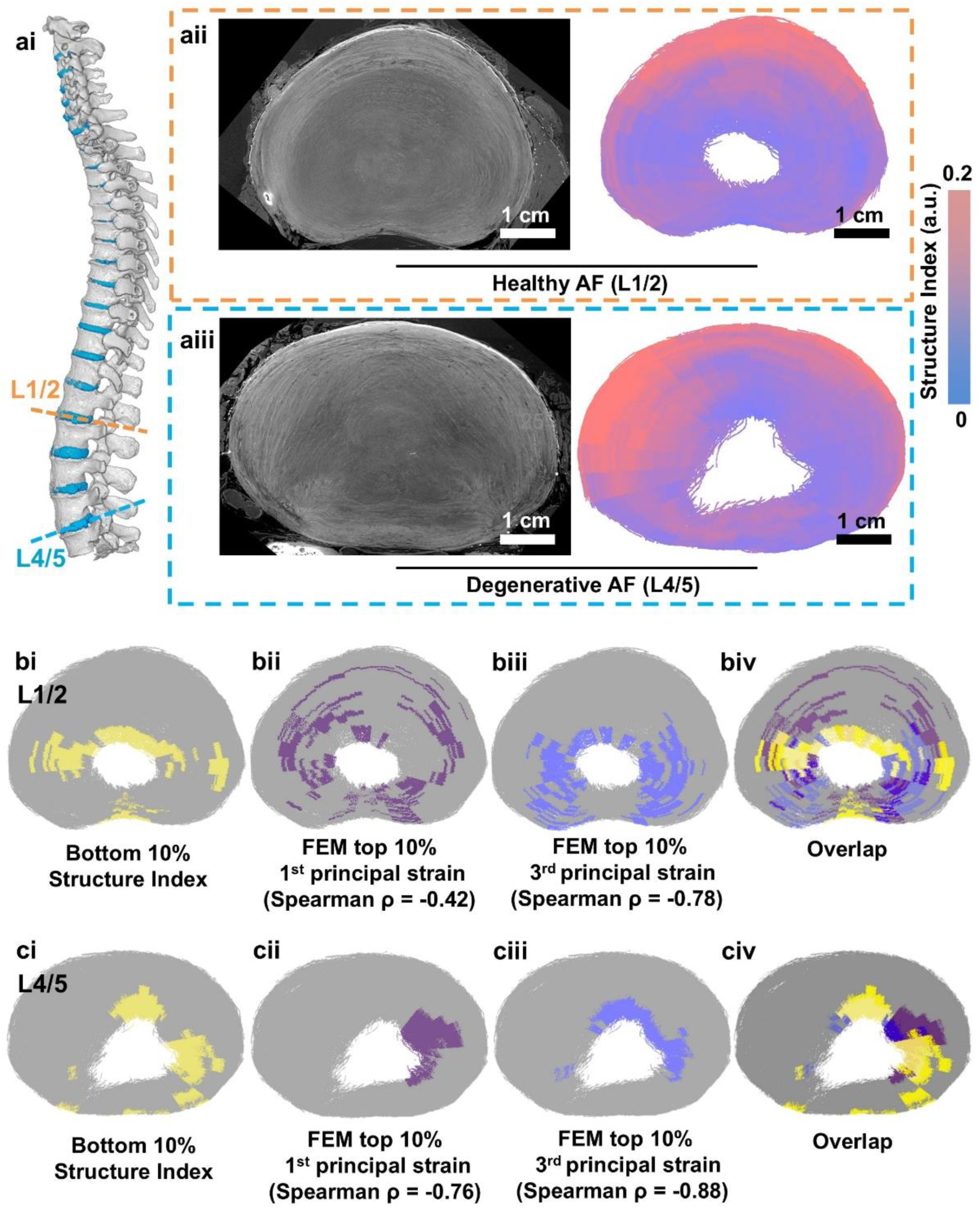
Structure Index mapping and high strain region prediction in both human healthy and degenerative discs. **(a)** Structural mapping of the AF in healthy (L1/2) and degenerative (L4/5) human IVDs; **(ai)** Anatomical location of analysed spinal segments; **(aii–aiii)** sCT images (left) and corresponding SI maps (right). The healthy AF **(aii)** exhibits a relatively symmetric spatial distribution, whereas the degenerative AF **(aiii)** shows marked redistribution of structural organisation; **(b)** Spatial correspondence between structurally predicted vulnerable regions and FEM-derived high-strain regions in the healthy disc; **(bi)** Bottom 10% SI regions; **(bii)** Top 10% first principal strain regions from FEM (Spearman ρ = −0.42); **(biii)** Top 10% third principal strain regions from FEM (Spearman ρ = −0.78); **(biv)** Spatial overlap between low SI and high-strain regions; **(c)** Corresponding analysis in the degenerative disc; **(ci)** Bottom 10% SI regions; **(cii)** Top 10% first principal strain regions from FEM (Spearman ρ = −0.76); **(ciii)** Top 10% third principal strain regions from FEM (Spearman ρ = −0.88); **(civ)** Spatial overlap between low SI and high-strain regions.

In contrast, the sCT image of degenerative human AF (L4/5) showed a more disrupted lamellar architecture, with less distinct nucleus pulposus–AF boundary (**Fig. 7aiii**). Correspondingly, SI analysis revealed a marked redistribution of low-SI regions. In the anterior AF, low-SI regions extended towards the outer two-thirds of the AF, whereas in the posterior–lateral AF they became restricted to the inner one-third (**Fig. 7aiii**). Compared with healthy discs, degeneration therefore produced more asymmetric SI distributions together with sharper local SI transitions. These findings indicate that degeneration disrupts the normal radial organisation of AF fibre architecture, shifting the disc from a relatively symmetric outer–inner structural pattern towards a polarised anterior–posterior mechanically vulnerable state. Based on the previously established SI–strain relationships across species, high-strain regions were predicted to occur in proximity to these low-SI domains. In healthy discs, these predicted vulnerable regions were distributed predominantly along the long axis of the disc geometry, whereas in degenerative discs they became increasingly polarised along the short axis.

To determine whether these structure-driven predictions corresponded to mechanically vulnerable regions, we next performed organ-scale FEM validation using the homogenised material properties derived from the multiscale framework. Regions corresponding to the bottom 10% SI exhibited similarity with FEM-predicted high-strain regions (**Fig. 7bi–biv**). In healthy discs, low-SI regions correlated with areas exhibiting the top 10% first and third principal strains, with Spearman correlations of −0.42 and −0.78, respectively. Importantly, this correspondence became stronger in degenerative discs, with correlations increasing to −0.76 for first principal strain and −0.88 for third principal strain (**Fig. 7ci–civ**).

These findings demonstrate that degeneration-driven structural reorganisation alters the spatial distribution of mechanically vulnerable regions within the human disc and that these vulnerable regions can be inferred directly from static structural organisation using the SI framework. The stronger SI–strain correspondence observed in degenerative discs further suggests that structural disorganisation is increasingly associated with mechanically vulnerable behaviour in degenerated tissue.

## Discussion

This study establishes a multiscale framework linking fibre architecture to mechanical response in the IVD. By combining sCT, fibre tracing, and digital volume correlation, we demonstrate that the AF fibre organisation is inversely associated to regions of high strain. The SI derived from structure tensor anisotropy and fibre orientation shows a robust negative correlation with principal strain magnitude across species. Importantly, this relationship was also observed across multiple strain measures, indicating that SI captures a broader architectural susceptibility to concentrated deformation. This structure–strain relationship enables prediction of mechanically vulnerable regions in human discs using static structural imaging alone. In addition to establishing a structure–strain relationship, this study provides a multiscale bridge linking fibril-level organisation to organ-scale mechanical behaviour. By integrating experimentally derived fibril morphology RVE homogenisation, we obtained effective anisotropic material properties that were subsequently applied in organ-scale FE models. This approach enables direct propagation of microscale structural information to macroscopic mechanical predictions^32^, overcoming a long-standing challenge in modelling fibre-reinforced biological tissues.

In terms of cross-species similarities and differences in AF architecture, our data indicate that structural conservation varies across hierarchical length scales. Gross disc and lamellar architectures retained broadly conserved organisational features across species^22,33,34^, despite species-specific differences in overall disc morphology and the spatial distribution of lamellae. Across species, the anterior AF consistently contained more lamellae than the posterior–lateral AF, while lamellar thickness generally decreased from the outer to inner AF. Together, these findings suggest that conservation at the lamellar scale lies primarily in shared regional and radial organisational principles rather than identical spatial distributions. In contrast, finer fibre-bundle organisation showed greater species- and region-dependent divergence. The outer AF generally exhibited more coherent fibre organisation, whereas the inner AF showed greater orientation dispersion and tortuosity, with θ-angle trends differing between species, particularly between mini-pig and rat or human discs. These hierarchical structural gradients may reflect adaptation to the distinct mechanical demands across the AF. The outer AF primarily resists circumferential tensile stresses generated by nucleus pulposus pressurisation, whereas the inner AF accommodates greater deformation and shear^12,19^. The progressive reduction in lamellar definition and increased fibre heterogeneity towards the inner AF may therefore facilitate local strain redistribution while maintaining overall disc integrity^31^.

Importantly, however, the relationship between structural organisation and mechanical response remained consistent across species. In both rat and mini-pig discs, lower SI values were preferentially associated with regions of elevated strain, indicating that the underlying mechanical principles governing fibre-reinforced disc behaviour are preserved despite morphological and architectural differences. The mechanical behaviour of the disc is strongly dependent on the integrity and organisation of its fibrous architecture, and its disruption alters local mechanical responses and mechanosensing^35^. Notably, the spatial overlap between the lowest 10% SI regions and the highest 10% strain regions was relatively limited, indicating that SI does not yield a direct point-by-point representation of instantaneous strain magnitude. Instead, it identifies structural configurations more susceptible to concentrated deformation. This interpretation is further supported by the progressive strengthening of structure–strain correlations as the analysis was restricted to increasingly high-strain domains. Within these regions, SI exhibited stronger correlations with strain than either structural anisotropy or fibre orientation alone, suggesting that the combined metric captures mechanically relevant architectural information more effectively than its individual components.

The capacity of SI to identify mechanically vulnerable regions enabled its application to human discs without mechanical loading. In healthy discs, the predicted vulnerable regions followed a relatively symmetric distribution along the long axis of the disc geometry. In contrast, degenerative discs exhibited a pronounced redistribution of low SI regions, resulting in asymmetric structural patterns. Specifically, low SI regions extended outward in the anterior AF but became confined to inner regions of the posterior–lateral AF. This redistribution suggests that disc degeneration disrupts the normal radial gradient of fibre organisation and leads to spatial reorganisation of mechanically vulnerable regions. Such changes are consistent with the recognised loss of tissue structural integrity and altered mechanical function in disc degeneration^36^. FEM provided an independent mechanical assessment of these predictions, with regions of low SI preferentially corresponding to regions of elevated strain. Moreover, the structure–strain association was stronger in more degenerated discs, implying that progressive structural disorganisation amplifies the local mechanical vulnerability.

These observations are consistent with previous studies showing that disc degeneration is associated with substantial alterations in the internal mechanical environment that are not captured by gross morphological changes alone ^12^. This highlights the importance of local tissue architecture, as quantified here by SI, in characterising degeneration-associated mechanical vulnerability. Experimental measurements using DVC have demonstrated elevated internal strains in degenerate discs, particularly in the lateral and posterior AF, together with increased tensile and shear strains near the endplates^37^. In addition, inward bulging of the inner annulus has been linked to local increases in compressive, tensile and shear deformation, which may predispose the tissue to circumferential tearing^37^. Structural failure of the disc has also been associated with pre-existing defects in the posterior AF and stress concentration at the annulus–endplate junction^38,39^, suggesting that local architectural heterogeneity plays a key role in initiating disc damage. Collectively, these findings support the interpretation that the redistribution of low SI regions reflects a spatial shift in mechanically vulnerable zones during degeneration, rather than a simple change in global structure. They also emphasise the need for mechanical models that can preserve the influence of local tissue architecture when evaluating such structure–mechanics relationships in human discs.

Addressing this need, our fibril-informed homogenisation framework provides a route for translating microscale organisation into organ-scale mechanics. Traditional FE models of the IVD often rely on simplified or assumed material properties^40,41^, which may not fully capture the influence of underlying fibre architecture. In contrast, our framework derives effective anisotropic properties from experimentally measured fibril geometry and spatial organisation before propagating these properties into the organ-scale model. This multiscale strategy therefore provides an independent mechanical basis for evaluating SI-predicted vulnerable regions in human discs, while reducing reliance on phenomenological material assumptions. More broadly, the correspondence between structurally vulnerable regions and mechanically concentrated deformation supports the concept that AF failure may preferentially emerge at locations where local fibre architecture is less able to sustain or redistribute load^42^.

Beyond the IVD, our framework may be applicable to other fibre-reinforced biological tissues. Many organs, including myocardium, tendon, and brain white matter, exhibit hierarchical fibre architectures that govern their mechanical behaviour. Structure-based metrics integrating anisotropy and fibre orientation, combined with multiscale homogenisation, may therefore provide a general strategy for linking microstructural organisation to organ-scale mechanics across biological systems.

Several limitations should be considered. The number of specimens analysed in each species was limited, and larger datasets will be necessary to further validate the observed cross-species relationships. DVC measurements were only performed in animal discs, and human strain distributions were inferred indirectly through FEM. Direct experimental measurement of internal strain in human IVDs would provide important validation of the predicted structure–mechanics relationships. In addition, the human specimens represented varying degrees of degeneration rather than healthy controls, limiting separation of species-specific architecture from degeneration remodelling. The SI is derived from local structural descriptors and does not explicitly incorporate long-range fibre connectivity, lamellar branching and merging, or interlamellar interactions^43^, all of which may contribute to load transfer. Its sensitivity to fibre number in poorly organised regions also indicates that sampling density should be considered when comparing SI across datasets. Furthermore, the structural information was derived from micron-resolution sCT of ex vivo specimens, and the extent to which these fibre-scale features can be recovered using clinically accessible modalities remains uncertain. Future studies should therefore combine larger human cohorts, direct mechanical validation and multimodal imaging to determine whether the structural signatures identified here can be approximated *in vivo* using MRI or clinical CT, ultimately enabling translation.

## Conclusion

This study provides a quantitative link between fibre architecture and mechanical response in the IVD. By establishing a cross-species structure–mechanics relationship and demonstrating its predictive capability in human discs, we show that mechanical vulnerability can be inferred directly from structural organisation. This work provides a foundation for structure-driven analysis of biomechanics in fibre-reinforced tissues. The concept of SI may provide a new biomarker for structure-based mechanical risk assessment in fibre-reinforced tissues, offering a potential route towards non-invasive risk assessment in diseases.

## Methodology

### Specimen preparation

The rat imaging datasets analysed in this study were reused from our previously published study^44^. In the original study, three male, 8-week-old Sprague Dawley rats used for this study were housed at the University of Manchester Biomedical Sciences Facility, Manchester, UK. Three lumbar segments were harvested from the three rats after euthanasia and frozen at −20℃ until use. The ethical approval for the use of experimental animals and tissue collection procedures was approved by the University of Manchester Animal Welfare and Ethics Review Board and followed the University’s Animal Research Policy and the UK Animal (Scientific Procedures) Act of 1986 under the Home Office Licence (#70/8858 or I045CA465). No additional experiments on rats were performed in the present study. Due to the sample availability, three rats each contributed one lumbar L2/3, L4/5, or L5/6 for this study.

Three female, 13- to 16-month-old Göttingen mini-pigs used for this study were obtained from Ellegaard Göttingen Minipigs A/S (Dalmose, Denmark). Three lumbar segments were harvested from the three mini-pigs after euthanasia and frozen at −20℃ until use. All the procedures were approved by the Local Animal Care Committee of Lithuanian University of Health Sciences (IACUC approval number: 2022-11-09-Nr.-G-2-224) and conducted in accordance with national and EU regulations on animal research. Three mini-pigs each contributed one lumbar L4/5 for this study.

Three 62- to 99-year-old male and female donors contributed three human spine segments^29^. The specimens were collected and then fixed in 4% formalin at the Laboratoire d’ Anatomie des Alpes Françaises (LADAF), Grenoble, France. All the procedures were approved by the Comité d’Éthique Scientifique et Pédagogique (CESP), Université Grenoble Alpes (#20-2023), and all specimens were de-identified. For all of the statistical analyses on human specimens, one L4/5 segment from each donor (donor 1 – 3) were utilised. For SI mapping, L1/2 and L4/5 from donor 2 were utilised as a comparison of ‘healthy’ and ‘osteoarthritis’ conditions, respectively. See **Table S2** for donor details.

Surrounding soft tissues of all specimens were carefully removed. Animal specimens were wrapped with polyethylene film to minimise dehydration during imaging. After that, they were mounted in a custom sample holder compatible with Deben CT500 or CT5000 rig. Human specimens were mounted in jars with Agar cubes on a high-precision rotation stage, and the HiP-CT protocol was followed^45^.

### *In situ* compression

For the rat specimens, a 1 N preload, equivalent to approximately 40% of the animal’s body weight, was first applied. This was followed by a 40 µm displacement step at a constant remote displacement rate of 0.1 mm/min. The specimens were then allowed to undergo 20 min of stress relaxation before imaging. Two scans were acquired for each specimen, corresponding to the preload and loading states.

For the mini-pig specimens, a 200 N preload, equivalent to approximately 80% of the animal’s body weight, was applied, followed by a 300 µm displacement step at a constant remote displacement rate of 0.5 mm/min. The specimens were then allowed to undergo 40 min of stress relaxation before imaging. Two scans were acquired for each specimen, corresponding to the preload and loading states.

No mechanical load was applied to the human specimens during imaging.

### synchrotron computed tomography imaging

sCT imaging was performed at beamline I13-2 of Diamond Light Source for the rat specimens and at beamline BM18 of the European Synchrotron Radiation Facility (ESRF) for the mini-pig and human specimens. The detailed acquisition parameters are summarised in **Table S3**.

Briefly, for the rat specimens, sCT imaging at Diamond Light Source was performed using a pink beam with a mean energy of 27 keV and an isotropic voxel size of 1.625 µm. Each scan took approximately 3 min and consisted of 1,800 projections acquired using fly scanning, with an exposure time of 0.1 s per projection. Tomographic reconstruction was performed using Savu.

For the mini-pig specimens, sCT imaging at ESRF BM18 was performed using a parallel, filtered polychromatic X-ray beam with a mean energy of 110 keV and an isotropic voxel size of 10.01 µm. The source-to-sample distance was 177 m and the propagation distance was 10 m. Each scan took approximately 1.7 min and consisted of 6,000 projections acquired with an exposure time of 0.015 s per projection.

For the human specimens, two HiP-CT acquisition configurations were used. Donor 1 specimen was imaged at a mean energy of 118 keV with an isotropic voxel size of 16.26 µm, a propagation distance of 32.3 m, 15,000 projections, and an exposure time of 0.008 s per projection. Donor 2 and 3 specimens were imaged at a mean energy of 90 keV with an isotropic voxel size of 20.19 µm, a propagation distance of 10.0 m, 12,000 projections, and an exposure time of 0.012 s per projection.

The mini-pig and human sCT datasets were reconstructed using the ESRF BM18 NightRail workflow^46^. Phase retrieval was performed using the fast Eikonal Phase Retrieval/WKB algorithm, which incorporates second-order eikonal transport corrections, polychromatic modelling, and local or non-local solvers depending on the refraction-shift regime. Ring artefact correction was subsequently applied, followed by filtered back-projection for tomographic reconstruction.

### Spatial mapping for local AF lamellar count

Bespoke MATLAB codes were developed to quantify the spatial distribution of AF lamellar number. Reconstructed TIFF image stacks and pre-prepared 3D AF segmentation masks were imported for analysis. The outer and inner boundaries of AF masks was identified, with the centroid of the inner boundary used as the geometric centre. Radial sampling lines were generated from the geometric centre across the AF inner and outer AF boundaries, with a pixel-wise circumferential spacing. Each valid profile was normalised and resampled into 256 equally spaced positions across the AF width. For each z slice, intensity profiles were sampled using a three-pixel-wide radial band. Profiles were smoothed, baseline-corrected and normalised before local intensity peaks were detected as lamellar features. The number of detected peaks was calculated within a 25-bin sliding window.

The analysis was repeated through the image z direction and normalised to 256 height positions, generating a 256 × 256 heatmap of local lamella counts across normalised AF width and height. A Gaussian filter was applied for visualisation, while the unsmoothed data was used for quantitative analysis.

### Fibre tracing and point cloud generation

Reconstructed sCT volumes were imported into Avizo 3D (Thermo Fisher Scientific, v.2024.1) for fibre tracing. The IVDs were first segmented to isolate AFs from surrounding vertebral bone and background using masking. The segmented AF regions were used for fibre tracing. The Cylinder Correlation module was firstly used to generate correlation and orientation fields representing locally coherent fibre alignment. The fibre tracts were subsequently reconstructed using the Trace Correlation Lines module. Tracing parameters (minimum seed correlation 85, minimum continuation 45, direction coefficient 0.2, minimum distance 5 voxels, minimum length 300 voxels, search-cone length 30 voxels, search-cone angle 35°, minimum step size 10%) were selected to capture coherent fibre-aligned structures while suppressing low-confidence features. The resulting fibre line sets were regularised using the Smooth Line Set module (smoothing coefficient 0.5, adherence to original data 0.25, 10 iterations) to achieve approximately uniform inter-point spacing (∼12 voxels) along each tract. The information of segments defined by adjacent point pairs, the coordinates of the points, and tortuosity were extracted into spreadsheets for downstream analyses. The local tensor components were calculated using bespoke MATLAB code. For each fibre location, a local radial direction was defined as the vector pointing from the fibre point towards the manually selected IVD centre. The tensor component along this local radial direction was referred to as the local radial tensor component.

### Digital volume correlation

DVC was performed using the open-source iDVC application (https://tomographicimaging.github.io/iDVC/index.html) to quantify strain fields in loaded animal IVDs. The preload sCT volume was used as the reference volume, and the corresponding loaded volume was used as the correlate volume. The identical point clouds and masks from previous fibre tracing workflow were used. Correlation subvolumes were defined as cubic subsets with a size of 20 voxels centred on each point-cloud location (**Fig. S9a**) Determination of optimal subvolume size was detailed in **Supplementary Methods** and **Fig. S10**. DVC was performed on 10,000 sampled points per subvolume with an estimated maximum displacement of 15 voxels. The zero-normalised sum of squared differences (ZNSSD) was used as the objective function, with tricubic interpolation applied for subvoxel intensity estimation. Twelve optimisation parameters were used for displacement fitting, and single-run mode was applied for each run. The resulting displacement fields were used to calculate Lagrangian strain tensors using the iDVC strain calculation routine. Local displacement gradients were determined using a strain window of 25 neighbouring points, from which the six strain tensor components were calculated. The first and third principal strains were subsequently obtained. These strain measurements were subsequently mapped onto the corresponding SI datasets for structure–strain correlation analysis (**Fig. S9b**). Strain accuracy and precision were determined from zero-strain DVC analysis as the mean absolute error and standard deviation of error, respectively, across the six strain tensor components. The resulting strain accuracy and precision were 0.030% and 0.043%, respectively.

### Structure Index calculation

Bespoke MATLAB codes were applied to calculate SI. Briefly, preload sCT volumes were loaded, followed by structure tensor computation at each point location from the fibre tracing-derived point cloud. For each point, a local subvolume (20 × 20 × 20 voxels) centred at the point coordinate was extracted from the image volume. This ensured that the structure tensor and strain tensor were registered at identical spatial locations and scale. Image greyscale intensity gradients within this subvolume were computed along the three spatial axes using MATLAB imgradientxyz function, and the local structure tensor J was formed from second-order gradient products averaged over the subvolume (**Equation 1**).

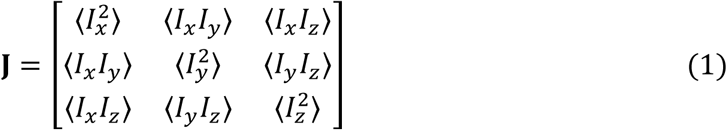

where *I_x_*, *I_y_*, *I_z_* are local image intensity gradients and ⟨⋅⟩ is averaging over the subvolume. The tensor was normalised by its trace and eigen-decomposed to obtain ordered eigenvalues *λ*_1_, *λ*_2_, and *λ*_3_ (*λ*_1_ ≥ *λ*_2_ ≥ *λ*_3_).

To normalise the depth of each point, inner and outer AF boundaries were estimated from the spatial distribution of fibre points in polar coordinates relative to the disc centre (manually selected) using a ray-based envelope reconstruction approach (**Fig. S11**). For each angular direction, the outer and inner AF boundaries were defined by the maximal and minimal radial distances of fibre points, respectively, followed by angular interpolation and smoothing to obtain continuous boundary curves. The radial depth *r* = [0,1], where *r* = 0 corresponds to the inner boundary and *r* = 1 to the outer boundary. Fibre θ orientation was obtained from the angle between fibre segment vectors and the disc axial (z axis).

We adapted and extended planarity and anisotropy measures derived from structure tensor analysis^47,48^ to define a SI using **Equation 2**.

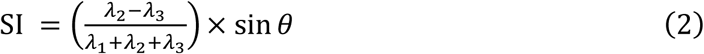

where *λ*_1_, *λ*_2_, *λ*_3_ are the eigenvalues of the structure tensor and *θ* is the fibre orientation relative to the disc axial. The first term quantifies the clarity of AF lamellar anisotropy, whereas the sin *θ* term weights the circumferential component of fibre orientation relative to the disc axial. The resulting Structural Index and depth values were exported for subsequent spatial correlation assessment.

### Structure–strain correlation analysis

Bespoke MATLAB codes were applied to quantify structure–strain correlation. For each point, lamellar anisotropy term, sin *θ* term, Structural Index, first and third principal strain magnitude were extracted. After that, the whole AF was partitioned into 36 circumferential sectors (10° each), and each circumferential sector was further subdivided into 36 radial depth bins based on sector-wise quantiles. Each point was assigned to its corresponding cell in this 36 × 36 grid according to spatial coordinates, and cell-wise mean values of Structural Index, its component terms, and principal strain magnitudes were computed from all points within each grid cell. Structure–strain correlation was quantified using Spearman rank correlation at the grid level within paired percentile subsets defined by progressively lower lamellar anisotropy term, sin *θ* term, or Structural Index (bottom 100%, 75%, 50%, 25%, and 10%), respectively, and correspondingly higher strain magnitude (top 100%, 75%, 50%, 25%, and 10%).

### Finite element modelling of human annulus fibrosus

A FE model of the human IVD was constructed by integrating fibre tracing data with tissue masks derived from sCT (**Fig. S12**). AF point cloud coordinates and segment connectivity obtained from fibre tracing were reconstructed into a 1D fibre network using bespoke MATLAB codes. The reconstructed fibre network was exported as beam elements in Abaqus CAE input format (.inp) to represent AF. The AF matrix and nucleus pulposus geometries were generated from their respective segmentation masks. These masks were meshed using tetrahedral elements with a target edge length of 325 μm and exported as Abaqus input files. All component models were then imported into Abaqus/Standard and assembled into a single FE model. Material properties were assigned to each component as listed in **Table 2 and 3.** Two reference points were then created to apply compressive loading. The superior reference point (RF1) was positioned above the centre of the disc, and the inferior reference point (RF2) was positioned below the disc centre. The superior and inferior disc surfaces were kinematically coupled to RF1 and RF2, respectively. The surface between AF matrix and nucleus pulposus was connected using surface-to-surface contact interaction with hard normal contact and isotropic tangential friction (friction coefficient = 0.02), allowing separation after contact. The reconstructed AF fibre network (embedded region) was embedded within the AF matrix (host) using the embedded region constraint. Axial compression along Z axis was applied by imposing a 300 μm displacement (equivalent to 5% remote strain) on RF2 while RF1 was encastred. The analysis was conducted using the implicit solver. After that, the first and third principal strains were extracted at nodal locations of the AF fibre network and used for subsequent structure–strain correlation analysis.

**Table 3.**
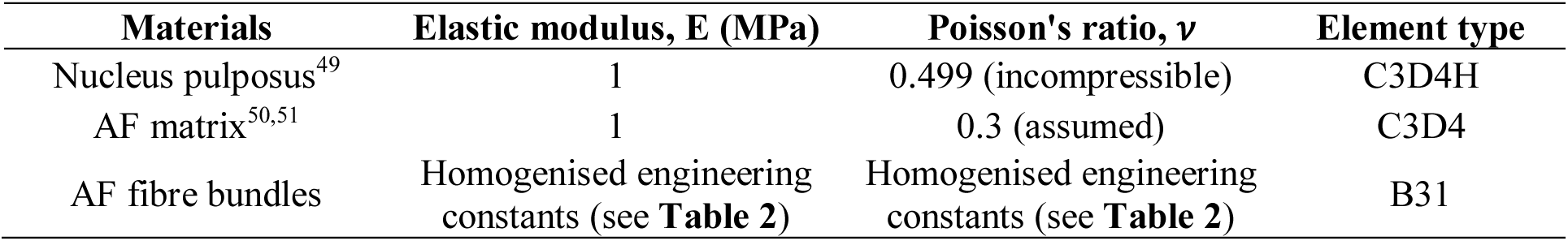
Material properties of the finite element model of rat lumbar spine segments according to the references with minor modifications.

| Materials | Elastic modulus, E (MPa) | Poisson's ratio, *ν* | Element type |
|---|---|---|---|
| Nucleus pulposus<sup>49</sup> | 1 | 0.499 (incompressible) | C3D4H |
| AF matrix<sup>50,51</sup> | 1 | 0.3 (assumed) | C3D4 |
| AF fibre bundles | Homogenised engineering constants (see **Table 2**) | Homogenised engineering constants (see **Table 2**) | B31 |

### Statistical analysis

All statistical analyses were performed using MATLAB unless otherwise stated. Data are presented as mean ± standard deviation unless indicated otherwise. Differences between groups were assessed using one-way analysis of variance (ANOVA) with Tukey’s post hoc comparison. Statistical significance was defined as P < 0.05. For fibril spatial organisation, the distribution of nearest neighbour distances was assessed for normality using the Kolmogorov–Smirnov test. Structure–strain relationships were quantified using Spearman rank correlation to account for potential non-linear and non-parametric associations. Correlations were computed at the grid level based on spatially averaged values. All statistical tests were two-sided.

## Data Availability Statement

Example sCT scan data of rat are available from the UCL Research Data Repository (DOI: 10.5522/04/26789212.v1). Example sCT scan data of mini-pig and human will be deposited in the HOAHub repository (https://human-organ-atlas.esrf.fr/) and made publicly available upon publication of the peer-reviewed article. All of the MATLAB codes used in this manuscript are available on Zenodo (DOI: 10.5281/zenodo.22111873).

## Acknowledgements

We sincerely thank Prof Michael J Sherratt for providing the experimental animals for this study. This work was supported by grants EP/V011235/1, EP/V011006/1, and EP/V011383/1 from the UK-Engineering and Physical Sciences Research Council, MR/V033506/1 from the UK-Medical Research Council, grant CZIF2021-006424 from the Chan Zuckerberg Initiative Foundation, and grant 2022-316777 from the Chan Zuckerberg Initiative DAF, an advised fund of Silicon Valley Community Foundation. We gratefully acknowledge Beamtime SM29784 at Beamline I13-2, Diamond Light Source; and Beamtime MD-1290 and MD-1389 at Beamline BM18, European Synchrotron Radiation Facility.

## Ethics approval statement

The ethics for using the rats for experiment and tissue collection procedures was approved by the University of Manchester Animal Welfare and Ethics Review Board and followed the University’s Animal Research Policy and the UK Animal (Scientific Procedures) Act of 1986 under the Home Office Licence (#70/8858 or I045CA465). The ethics for using the mini-pigs for experiment and tissue collection procedures was approved by the State Food and Veterinary Service of Lithuania (#G2-224) and conducted in accordance with national and EU regulations on animal research. Human cadaveric specimens were obtained from the Grenoble-Alpes body donation centre (Université Grenoble Alpes, France) under institutional approval (reference numbers 55-2023, 69-2023, and 72-2023). The authors sincerely thank those who donated their bodies to science so that anatomical research could be performed. Results from such research can potentially increase humanity’s overall knowledge that can then improve patient care. Therefore, these donors and their families deserve our highest gratitude.

## Author Contributions

Conception and design of the study: P.D.L. and J.C. Acquisition of data: J.C., A.P., A.S., I.H., J.B., T.U., P.D.L. and H.D. Interpretation of data, revision of the manuscript, final approval and agreement to be accountable for all aspects of the work: all authors. Drafting of the manuscript: J.C., A.P., A.S., and P.D.L.

