## Supplementary Information for "Cross-species structure–mechanics prediction in the intervertebral disc"

### Supplementary Methods

#### Synthetic lamellar fibre phantom

A 3D synthetic phantom was constructed using bespoke MATLAB code to validate the behaviour of the Structure Index under controlled structural conditions. The phantom was defined within a 100 × 100 × 100 voxels cubic domain. Seven parallel layers were generated along the transverse (y) direction with uniform spacing. Within each layer, fibres were modelled as independent cylindrical structures with a diameter of 1.5 voxels and extended across the entire domain. For each fibre, a line passing through a layer-specific seed point was intersected with the cubic boundary, ensuring that both endpoints lay on opposite faces of the cube. To preserve a discrete lamellar architecture, fibre spacing within each layer was constrained to exceed twice the fibre diameter, preventing overlap and the formation of continuous sheet-like structures.

Three independent parameters were introduced to control the structural configuration of the phantom:

1) The degree of lamellar organisation was varied using a scalar parameter (0–1), which controlled both positional perturbation of fibres and deviation from a preferred orientation. At low values, fibres exhibited strong spatial and directional randomness, whereas at high values, fibres were confined to well-defined layers with minimal deviation;

2) Fibre orientation relative to the z axis was controlled by an angle θ (0°–90°). A preferred direction vector was defined for each configuration, and individual fibre orientations were generated as a weighted combination of this preferred direction and a random unit vector;

3) The number of fibres was varied to assess the sensitivity of the SI to fibre sampling. A series number of fibres (3, 6, 9, 12, and 15 fibres) was set to per layer. Lower fibre numbers were generated by random subsampling the full 15-fibre population, thereby varying the number of tracked fibres while preserving the overall fibre organisation.

The resulting binary volumes were smoothed using a three-dimensional Gaussian filter ($\sigma = 0.35$ voxels) to generate continuous intensity gradients. Structure tensors were computed from spatial intensity gradients, followed by Gaussian smoothing ($\sigma = 3$ voxels) to capture mesoscale organisation. Eigenvalues of the tensor ($\lambda_1 \geqslant \lambda_2 \geqslant \lambda_3$) were used to compute the Structure Index.

#### Determination of optimal subvolume size for iDVC

The optimal subvolume size for iDVC was determined using a repeated-scan strategy under zero-strain state. Two independently acquired grayscale volumes of the same specimen were analysed with identical iDVC configuration to quantify measurement noise in the absence of deformation. Cubic subvolumes with edge lengths ranging from 10 to 100 voxels were evaluated. For each subvolume size, displacement fields were computed using a fixed configuration: 10,000 sampling points, 12 degrees of freedom, zero-mean normalised sum of squared differences (ZNSSD) objective function, tricubic interpolation, and the maximum expected displacement was set to 1 voxel. Performance was assessed using the objective function minimum (Objmin) and displacement uncertainty (x, y, z components). The optimal subvolume size was selected as the smallest size at which (1) the Objmin reached a stable regime and (2) displacement uncertainty was minimised and plateaued.

**Homogenisation of collagen fibril mechanical properties**

To determine the equivalent engineering constants of collagen fibril networks in the AF, a multiscale homogenisation framework was established based on experimentally derived fibril morphology and spatial organisation. Transmission electron microscopy images of bovine AF from [1] were analysed to quantify collagen fibril geometry and spatial distribution. Image processing was performed through binarisation and boundary detection to identify individual fibrils and extract their geometrical features (**Figure S8a**). The nearest neighbour distance between fibril centres was further calculated to characterise spatial packing, which followed a normal distribution analysis using Kolmogorov–Smirnov test.

Based on the extracted distributions, two-dimensional RVEs were constructed using a stochastic packing strategy, in which circular inclusions representing collagen fibrils were randomly distributed within a square domain under non-overlapping and minimum distance constraints. Using bespoke MATLAB codes, a series of RVEs containing increasing numbers of fibrils (5–50) were generated to evaluate statistical representativeness. Each 2D RVE was then extruded along the out-of-plane direction to form a 3D RVE, preserving the in-plane spatial organisation of fibrils (**Figure S8b**).

FE homogenisation was performed in Abaqus CAE (v.2023, Simulia, Providence, US) using the EasyPBC plugin (v1.4), which enables the automated implementation of periodic boundary conditions (PBCs) for RVEs. The use of PBCs ensures deformation compatibility and stress continuity across opposite faces of the RVE, thereby minimising boundary effects and enabling accurate estimation of effective material properties. Both fibrils and surrounding matrix were modelled as linear elastic materials as shown in **Table S1**. Perfect bonding between the two phases was assumed.

To determine the minimum fibril number, a convergence analysis was conducted by evaluating the percentage change in the volume-averaged von Mises strain as a function of fibril number. To determine the minimum RVE mesh size, a convergence analysis was conducted by evaluating the percentage change in the volume-averaged first principal strain as a function of mesh size (50–500 nm). A convergence threshold of 1% was adopted, and the corresponding RVE mesh size was selected for subsequent homogenisation.

For each RVE, periodic boundary conditions were applied in all three directions, and a set of independent loading cases was automatically generated using EasyPBC. These included three normal strain loadings along the principal directions and three shear loadings to fully characterise the orthotropic mechanical response. The apparent stress–strain response of the RVE was obtained from volume-averaged fields computed over the entire domain.

Using the RVE, the effective elastic constants, including $E_{11}$, $E_{22}$, $E_{33}$, $G_{12}$, $G_{13}$, $G_{23}$, and Poisson's ratios, were extracted directly from the homogenisation outputs of EasyPBC. The resulting homogenised properties were assembled into an orthotropic constitutive model representing the equivalent behaviour of the fibril–matrix composite. These equivalent material parameters were subsequently assigned to the AF fibre regions in the macroscale finite element model, enabling the integration of microscale fibril organisation into organ-scale mechanical simulations.

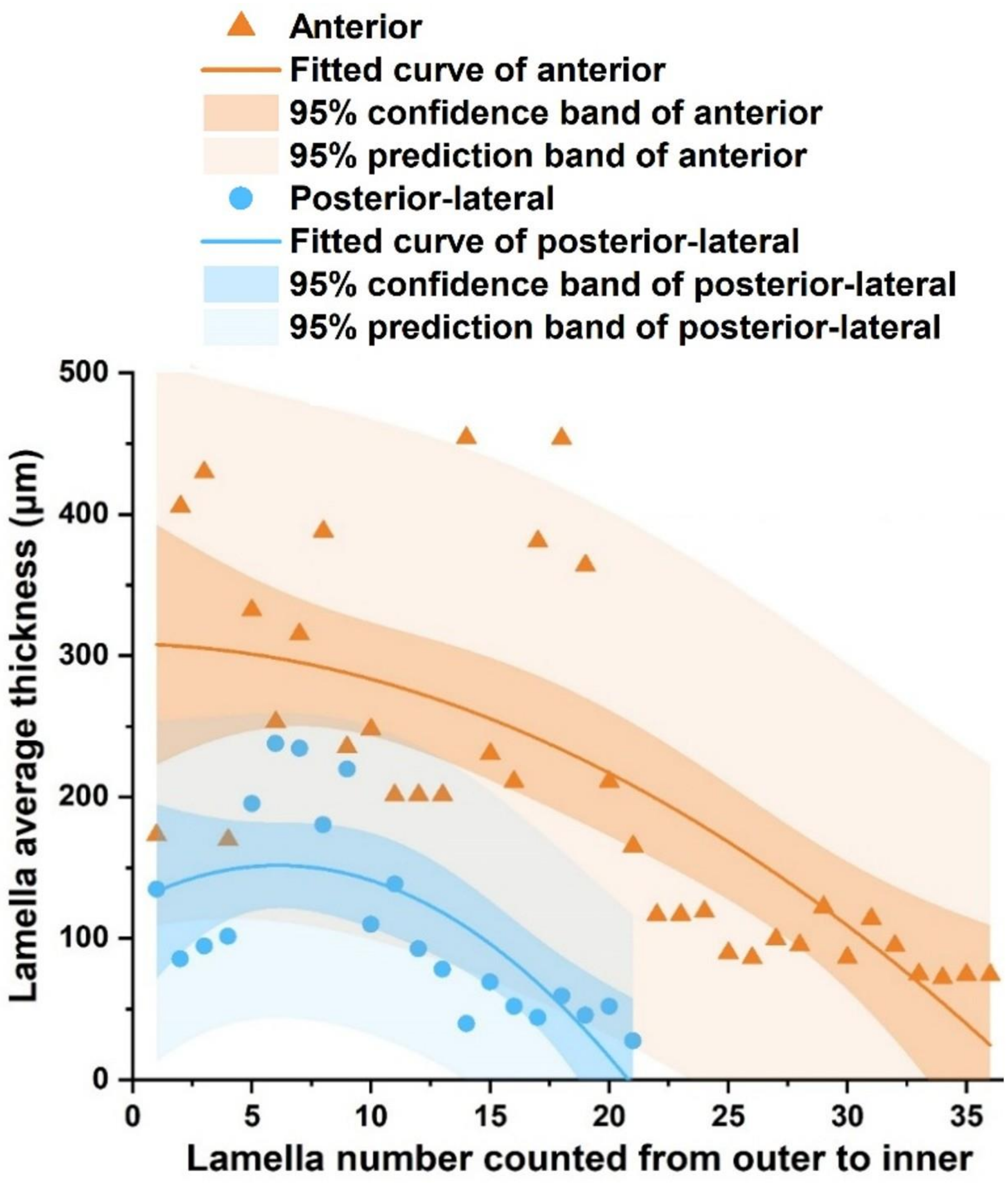


**Figure S1. Analysis of depth-dependant lamellar thickness in a representative mini-pig.** Lamellar thickness as a function of radial position from outer to inner AF, with fitted curves and confidence intervals for anterior and posterior–lateral regions;

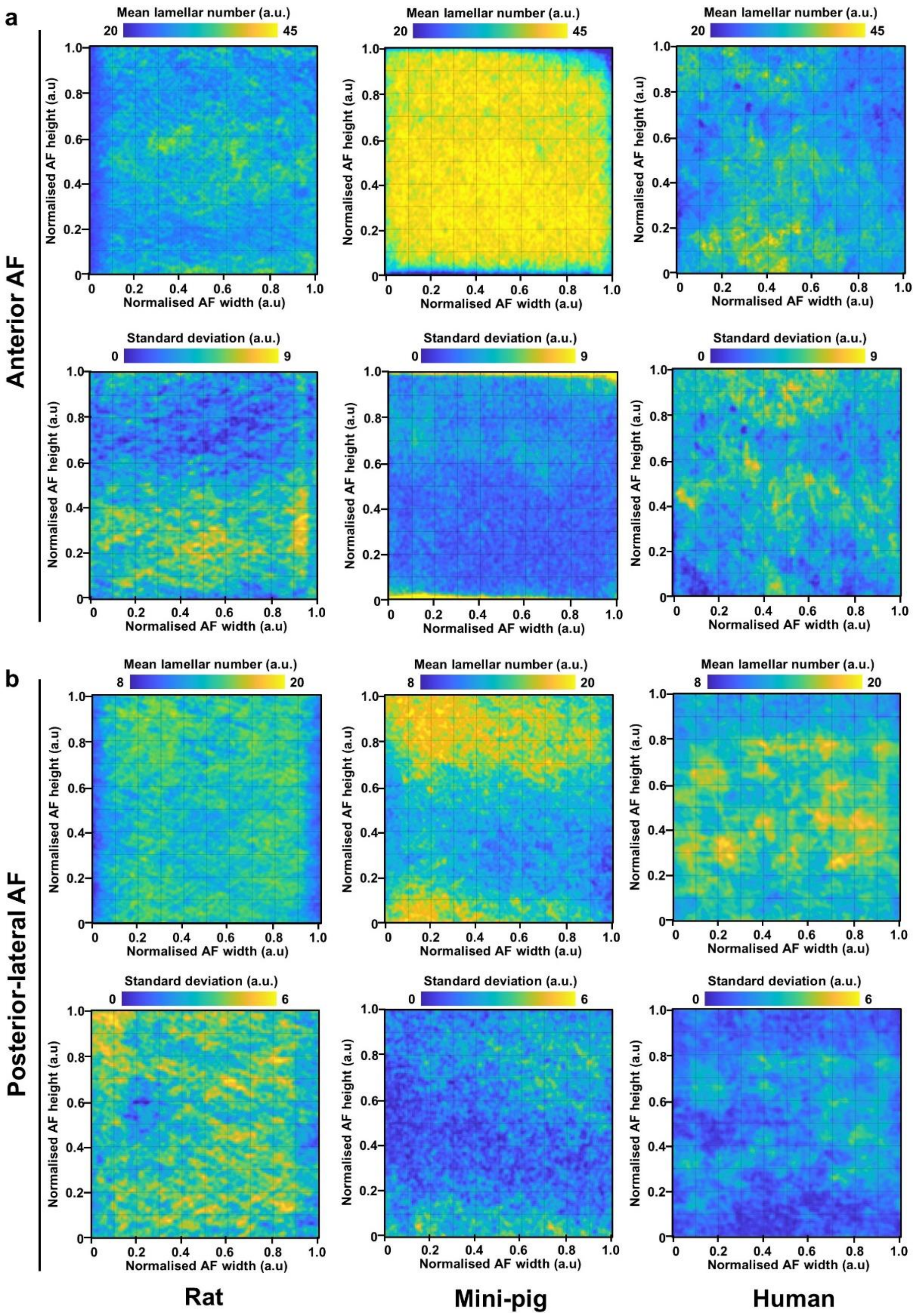


**Figure S2. Spatial distribution and variability of AF lamellar number across species.** Mean lamellar number and corresponding standard deviation (bottom) across normalised AF width and height in the **(a)** anterior or **(b)** posterior–lateral AF of rat, mini-pig, and human IVDs.

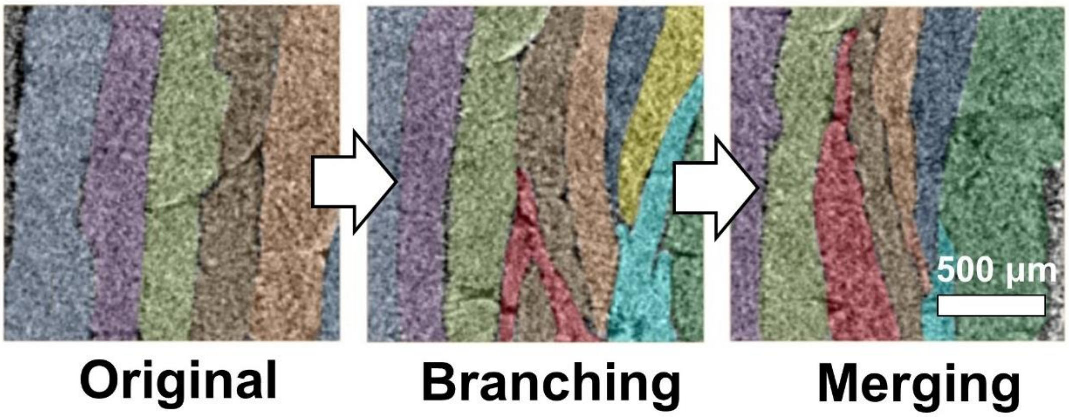


**Figure S3. Representative sCT images illustrating lamellar branching and merging within the AF.** Individual lamellae are pseudo-coloured to highlight their trajectories. A lamella branches (red) into adjacent layers and subsequently merges with neighbouring lamellae.

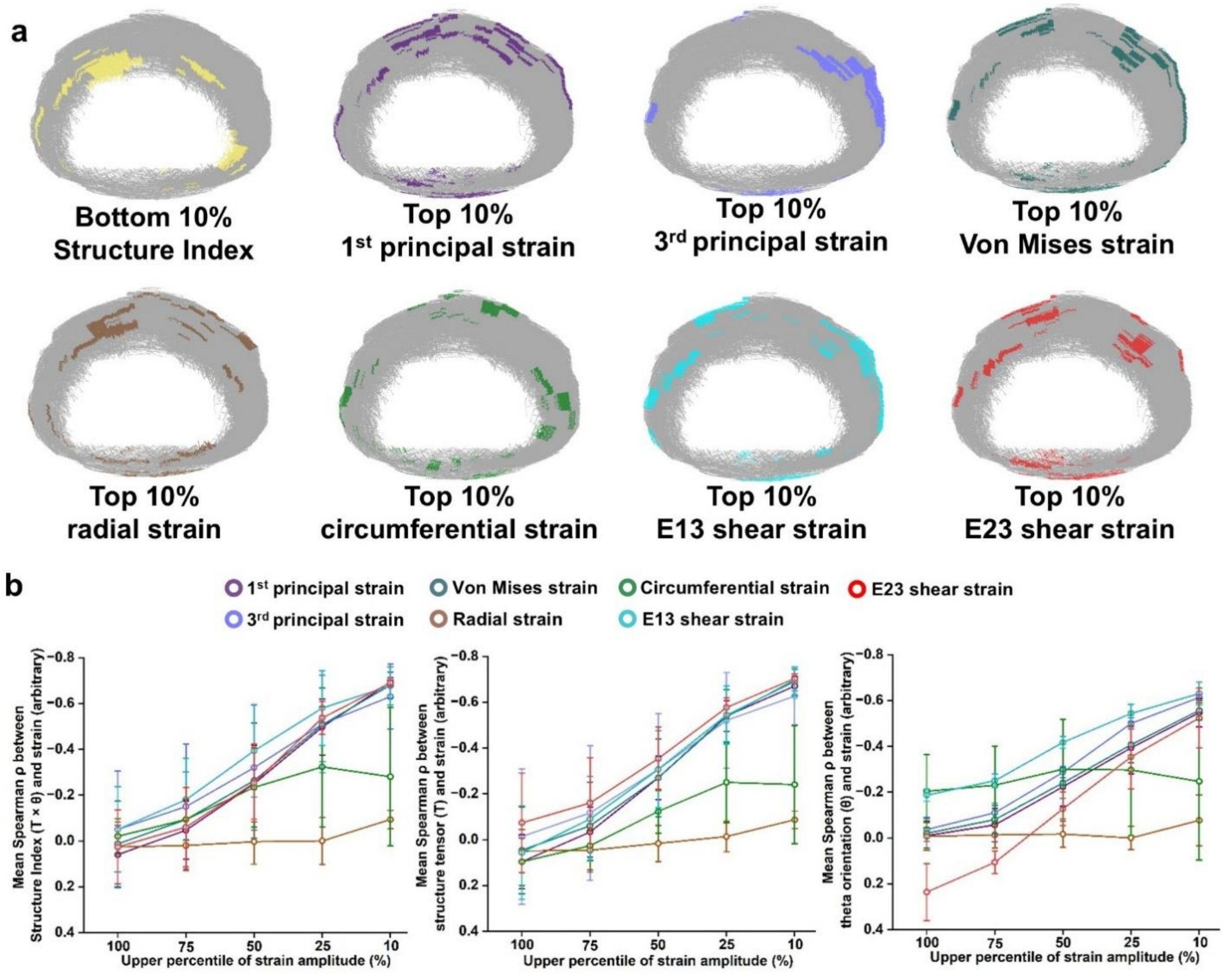


**Figure S4. Spatial association between Structure Index and strain fields in rat intervertebral discs. (a)** Spatial distributions of the bottom 10% SI values and the top 10% of first and third principal, Von Mises, radial, circumferential, E13 shear and E23 shear strains. Highlighted regions indicate the corresponding low-SI or high-strain domains within the AF; **(b)** Spearman correlations between SI and each strain measure calculated after progressively restricting the analysis to the upper 100%, 75%, 50%, 25% and 10% of strain magnitude.

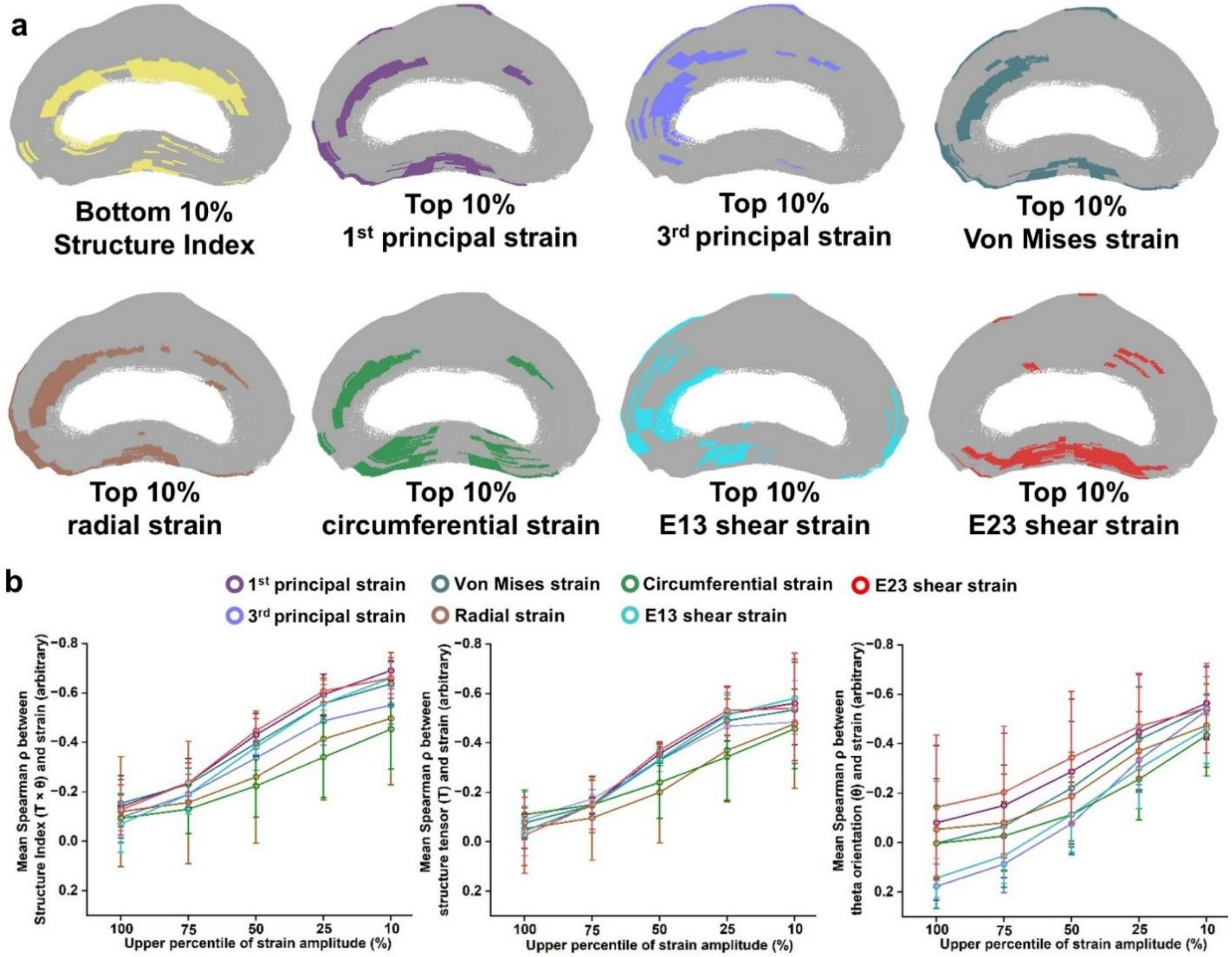


**Figure S5. Spatial association between Structure Index and strain fields in mini-pig intervertebral discs. (a)** Spatial distributions of the bottom 10% SI values and the top 10% of first and third principal, Von Mises, radial, circumferential, E13 shear and E23 shear strains. Highlighted regions indicate the corresponding low-SI or high-strain domains within the AF; **(b)** Spearman correlations between SI and each strain measure calculated after progressively restricting the analysis to the upper 100%, 75%, 50%, 25% and 10% of strain magnitude.

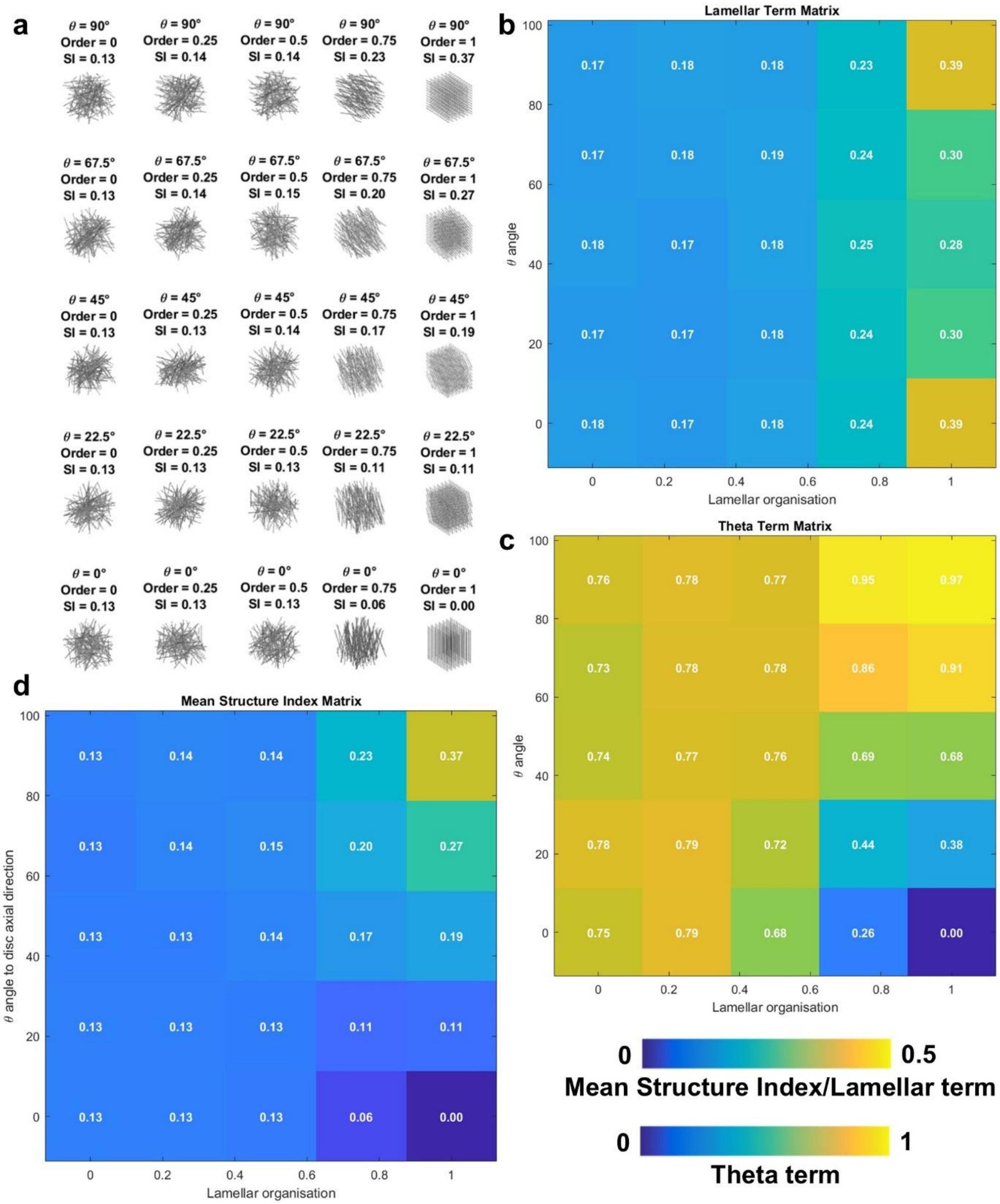


**Figure S6. Phantom validation of the Structure Index. (a)** Matrix of 3D synthetic lamellar fibre phantoms with independently controlled lamellar organisation and fibre orientation angle θ; **(b)** Lamellar term matrix; **(c)** Orientation term sin(θ) matrix; **(d)** Mean Structure Index matrix. SI = Structure Index.

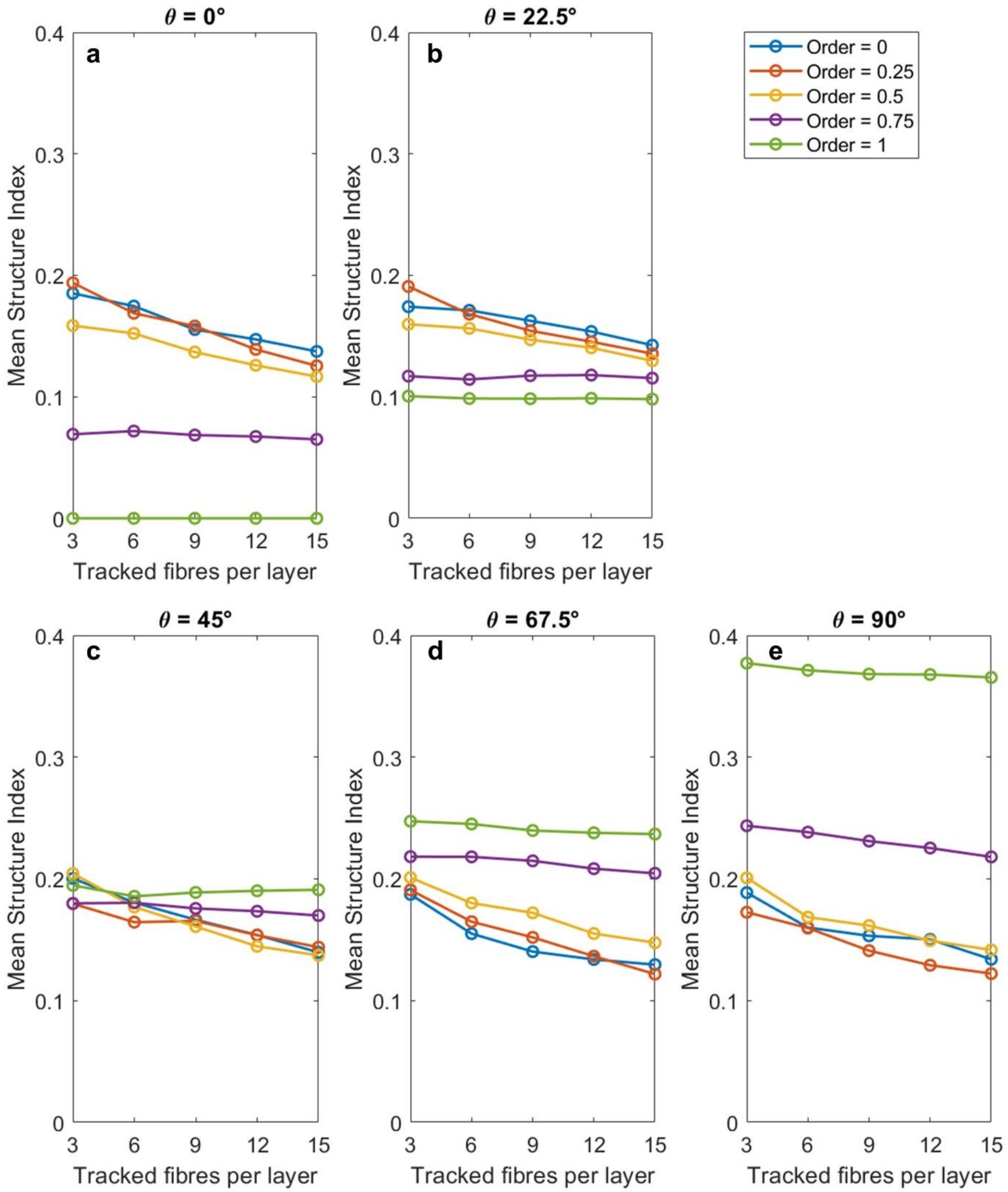


**Figure S7. Influence of fibre number on Structure Index across fibre orientations and organisation.** Mean SI as a function of the number of fibres per layer for fibre orientations of **(a)** θ = 0°, **(b)** 22.5°, **(c)** 45°, **(d)** 67.5°, and **(e)** 90°. Lines represent different levels of fibre organisation (Order = 0–1, from completely random to parallel within each layer).

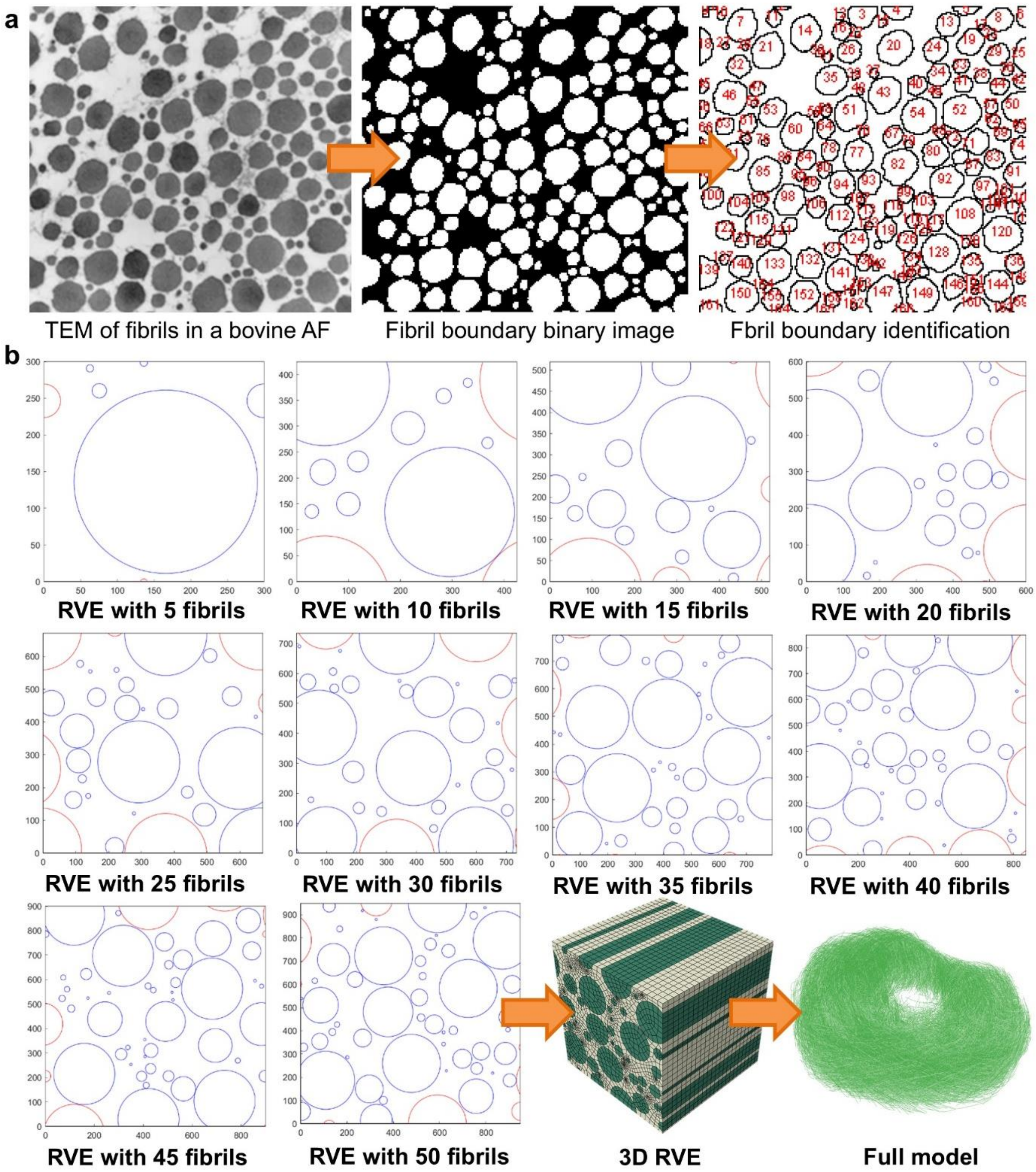


**Figure S8. From fibril morphology to RVE-based homogenisation of the AF. (a)** Collagen fibrils were segmented from transmission electron microscopy images of bovine AF[1]; **(b)** Stochastic RVEs were generated using the measured distributions. RVE size convergence was evaluated using increasing fibril numbers (5–50), followed by 3D extrusion and finite element homogenisation. The resulting effective properties were assigned to the full-scale mode. TEM: Transmission electron microscopy.

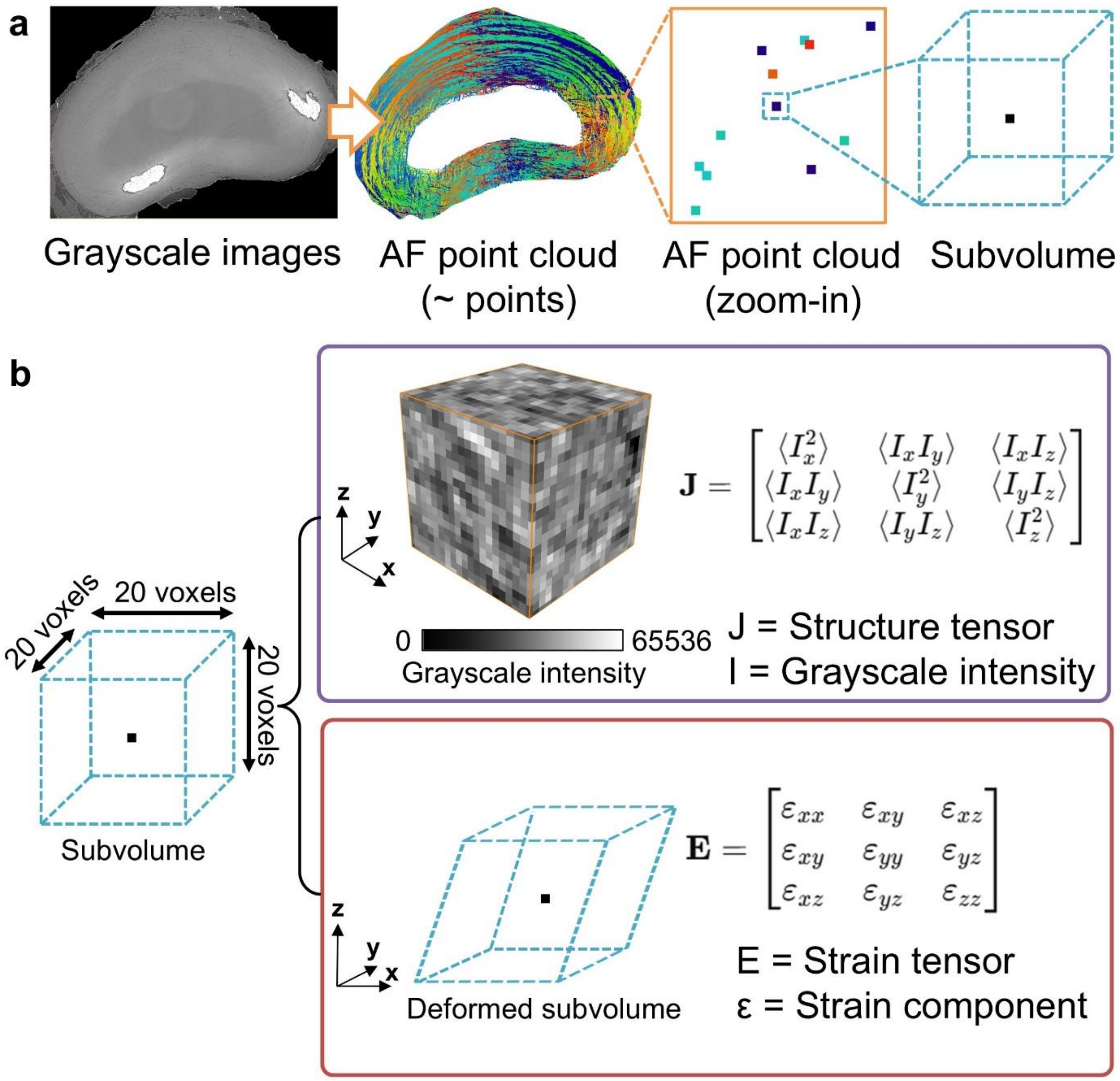


**Figure S9. Workflow for structure and strain tensor calculation from AF point cloud. (a)** AF point cloud generation; **(b)** Structure and strain tensor calculation.

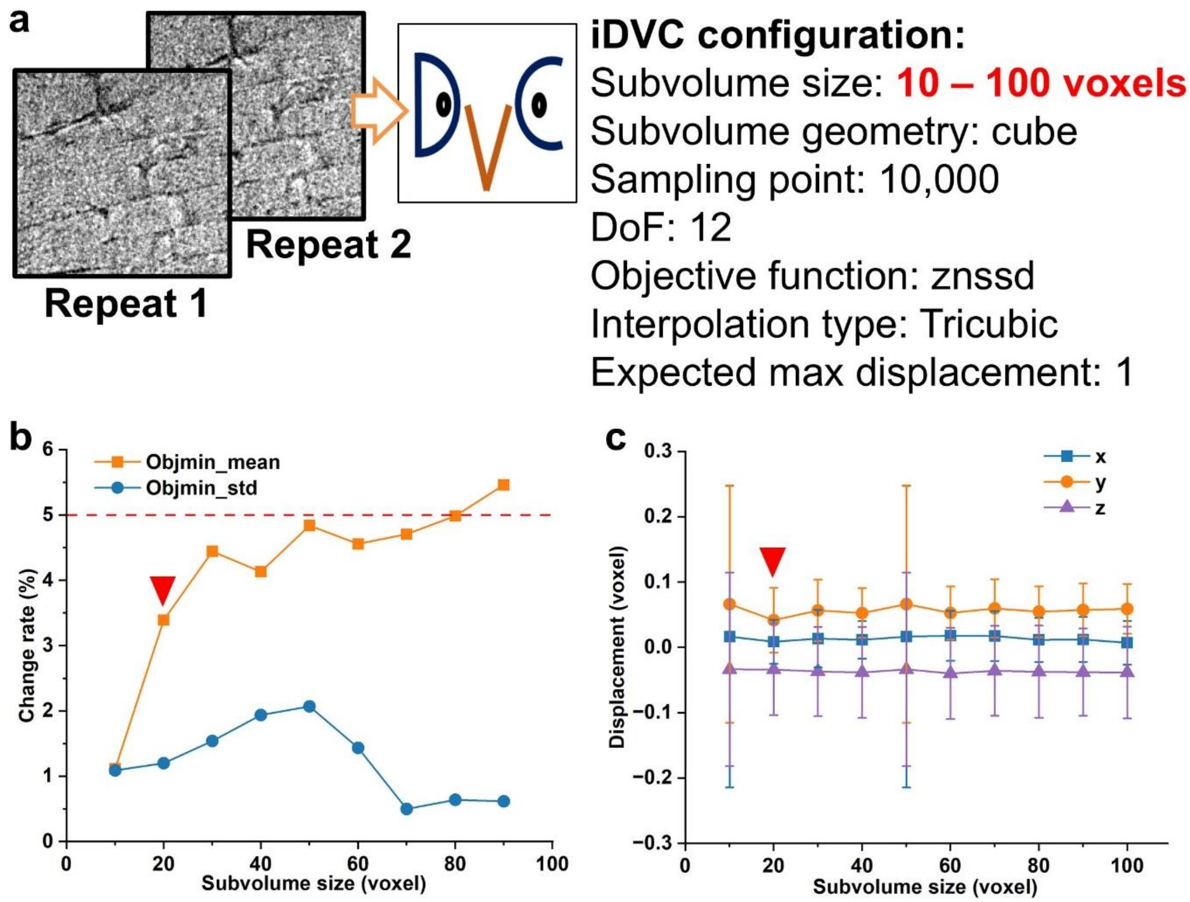


**Figure S10. Determination of optimal subvolume size for iDVC analysis. (a)** Schematic of the repeated scans used to evaluate iDVC performance under identical conditions and iDVC configuration. **(b)** Change rate of the objective function (Objmin) with subvolume size. The dashed line indicates the stability threshold, and the red marker indicates the selected optimal subvolume size. **(c)** Displacement uncertainty (x, y, z components) with subvolume size. The red marker indicates the selected optimal subvolume size.

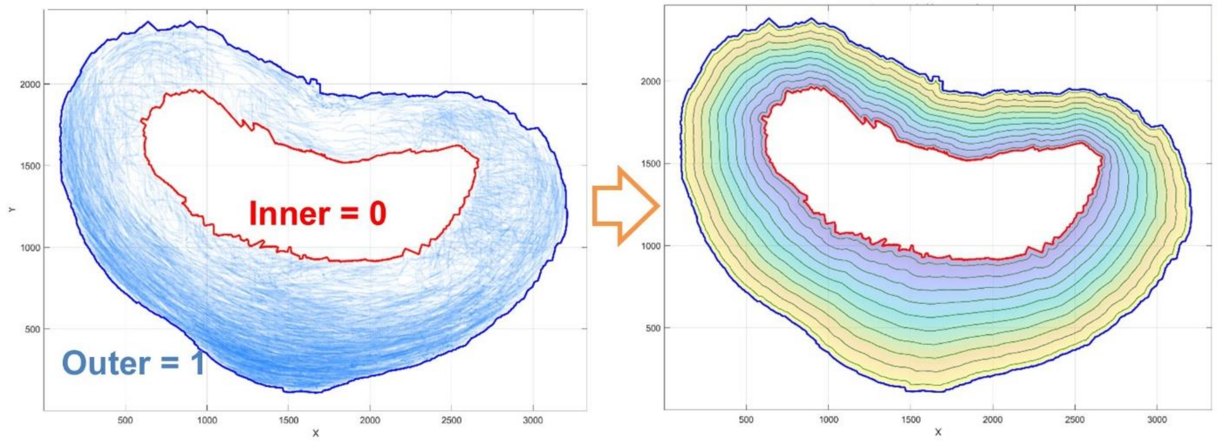


**Figure S11. Depth mapping of AF point cloud.**

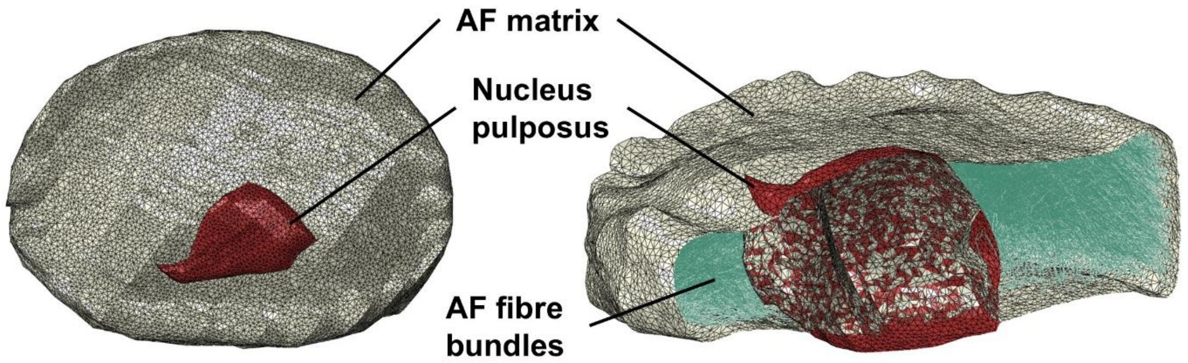


**Figure S12. FEM construction of the intervertebral disc.** 3D reconstruction of the intervertebral disc finite element model, including the AF matrix, embedded AF fibre bundles, and nucleus pulposus.

**Table S1.** Material properties of the collagen fibril RVEs.

| Materials | Elastic modulus, E (MPa) | Poisson's ratio, $\nu$ | Element type |
|---|---|---|---|
| Collagen fibril[2] | 350 | 0.3 (assumed) | C3D8R |
| Matrix[3,4] | 1 | 0.3 (assumed) | C3D8R |

**Table S2.** Information of human spine segment donors.

| Donor ID | Vertebral segment | Age | Sex | Ethnic group | Body height and weight | Dimension (L × W × D, cm) | Medical history |
|---|---|---|---|---|---|---|---|
| **1** | L3 – L5 | 81 | F | Caucasian | 160 cm, 55 kg | 12 × 8 | No known diseased |
| **2** | C1 – L5 | 62 | M | Caucasian | 175 cm, 70 kg | 62 × 13 × 17 | No known diseased |
| **3** | L1 – S1 | 99 | F | Caucasian | 165 cm, 65 kg | 23 × 12 × 11 | Lumbar osteoarthritis, sacroiliac osteoarthritis, left coxarthrosis, fracture of the right lesser trochanter, post-fracture left wrist algoneurodystrophy |

**Table S3.** sCT acquisition parameters for mini-pig and human specimens.

| **Sample** | Rat 1 – 3 | Mini-pig 1 | Mini-pig 2 | Mini-pig 3 | Human donor 1 | Human donor 2 | Human donor 3 |
|---|---|---|---|---|---|---|---|
| **Voxel Size** | 1.625 μm | 10.01 μm | | | 16.26 μm | 20.192 μm | |
| **Optic** | 2× objective lens; total magnification 4× | DZoom_X0.3574989539158418 | | | Fixed_X0.125 | DZoom_X0.3574989539158418 | |
| **Source** | I13-2 pink beam; source-to-sample distance 230 m | BM18 lateral beam 1.1 T (lateral shift by -10.0 mm at primary slits) | | | BM18 lateral beam 1.1 T (lateral shift by -15.0 mm at primary slits) | BM18 lateral beam 1.1 T (lateral shift by -17.3 mm at primary slits) | |
| **Filters** | Pyr. Graphite 0.28 mm + 1.06 mm, Al 3.2 mm, Fe 0.14 mm | sapphire 12.5 mm, SiO2 55 mm | | | sapphire 5mm, silver 0.3mm, molybdenum 0.23mm | sapphire 2.5mm, silver 0.1mm | |
| **Average Energy** | 27 keV | 110 keV | | | 118 keV | 90.0 keV | |
| **Surface Dose Rate** | N/A | 3.4 Gy/s | | | 15 Gy/s | 34 Gy/s | |
| **Integrated Dose to Sample** | 27 kGy | 1.4 kGy | | | 5.0 kGy | 11 kGy | |
| **Propagation Distance** | 0.5 m | 10.0 m | | | 32.3 m | 10.0 m | |
| **Scintillator** | 500 μm LuAG So14 | LuAG 2000 | | | LuAG2000 refl | GAGG1000 refl | |

| | | | | | | | |
|---|---|---|---|---|---|---|---|
| **Sensor** | pco.edge 5.5 camera | IRIS_15 (5056 × 600 pixels) | | | IRIS_15 (5056 × 600 pixels) | IRIS_15 (2528 × 512 pixels in 2 × 2 binning) | |
| **Single Projection Field of View** | 4.2 × 3.5 mm² | 50.61 × 10.25 mm² | | | 82.21 × 9.76 mm² | 51.05 × 10.34 mm² | |
| **Number of Projections** | 1800 | 6000 | | | 15000 | 12000 | |
| **Exposure Time** | 0.1 s | 0.015 s | | | 0.008 s | 0.012 s | |
| **Jar Mounting** | N/A | N/A | | | 145 mm diameter sample container immersed with 3 mm of 70% ethanol around it, held in 2 mm thick polycarbonate tube | 145 mm diameter sample container immersed with 3 mm of 70% ethanol around it, held in 2 mm thick polycarbonate tube | |
| **Horizontal Acquisition Scheme** | 180° fly-scan | Half-Acquisition | | | Half-Acquisition | Quarter-Acquisition | |
| **Vertical Acquisition Scheme** | Single scan | Single scan | | | Helical with 4.8 mm vertical displacement between scans | Z-series with 7.0 mm vertical displacement between scans | |
| **Reference Geometry** | Reference in air | Reference in air | | | Reference centered in equivalent container filled with PBS, full integration per turn | Reference centered in equivalent container filled with PBS, full integration per turn | |
| **Total Field of View Diameter** | N/A | 51.0 mm | 50.9 mm | 51.0 mm | 150.5 mm | 140.0 mm | |
| **Total Field of View Height** | N/A | 10.3 mm | | | 649.3 mm | 141.5 mm | 227.0 mm |
| **Number of Turns** | N/A | 1 | | | 133 | 2 × 20 | 2 × 32 |
| **Time per Turn** | N/A | 1.7 min | | | 2.2 min | 2.6 min | |
| **Total Time** | 3 min | 0.1 h | | | 4.9 h | 1.9 h | 2.8 h |
| **Reconstruction protocol** | Savu; flat- and dark-field correction; lens distortion correction; ring artefact removal; filtered back-projection with Paganin phase retrieval (δ/β = 100) | Processing with Nightrail. Tomographic reconstruction using filtered back-projection with single distance phase retrieval coupled with unsharp mask | | | Processing with Nightrail. Ring corrections by partial average of projections over several turns, tomographic reconstruction using filtered back-projection with single distance phase retrieval | Processing with Nightrail. Ring corrections by partial average of projections over several turns, tomographic reconstruction using filtered back-projection with single distance phase retrieval coupled with unsharp mask | |

| | | | | |
|---|---|---|---|---|
| | | | coupled with unsharp mask | |
| **Post-Processing** | 16-bit TIFF stacks | 16 bit conversion, binning with increasing level until reaching 8 GB | 16 bit conversion, jp2k compression, binning with increasing level until reaching 8 GB | 16 bit conversion, jp2k compression, binning with increasing level until reaching 8 GB |
